# Role of PLK1 in the epigenetic maintenance of centromeres

**DOI:** 10.1101/2024.02.23.581696

**Authors:** Duccio Conti, Arianna Esposito Verza, Marion E. Pesenti, Verena Cmentowski, Ingrid R. Vetter, Dongqing Pan, Andrea Musacchio

## Abstract

The centromere, a chromosome locus defined by the histone H3-like protein CENP-A, seeds the kinetochore to bind microtubules during cell division. Centromere maintenance requires CENP-A to be actively replenished by dedicated protein machinery in the early G1 cell-cycle phase, compensating for its two-fold dilution following DNA replication. Cyclin-dependent kinases (CDKs) limit CENP-A deposition to once per cell cycle and function as negative regulators outside early G1. Antithetically, Polo-like kinase 1 (PLK1) promotes CENP-A deposition in early G1, but the molecular details are still unknown. We reveal a phosphorylation network that recruits PLK1 to the deposition machinery to control a conformational switch required for licensing the CENP-A deposition reaction. Our findings solve the long-standing question of how PLK1 contributes to the epigenetic maintenance of centromeres.

**One-Sentence Summary:** PLK1 licenses epigenetic maintenance of centromeres by regulating a conformational switch on the MIS18α protein.

## Introduction

During cell division, kinetochores choreograph the segregation of sister chromatids by mediating their interaction with the mitotic spindle (*1*). Kinetochores crop up from specialised chromatin loci called the centromere (*2–6*). The maintenance of centromeres’ position along the chromosome axis is of utmost importance for cell viability (*7*). With few exceptions, the specialised histone Centromeric Protein A (CENP-A) defines the location of centromeres, functioning as an epigenetic marker (*8–10*). In higher eukaryotes, CENP-A dynamics on chromatin differ from those of canonical H3.1 histones, which are incorporated during DNA replication (*11, 12*). CENP-A replenishment in telophase/early G1 phase compensates for its two-fold dilution during S-phase (*13, 14*) (**Fig. 1A**). A dedicated group of conserved proteins, collectively termed the CENP-A deposition machinery, orchestrates this unique behaviour. Specifically, Holliday junction recognition protein (HJURP), a CENP-A-specific chaperone, stabilises the cytosolic form of CENP-A (*15, 16*). Targeting HJURP to centromeres requires the octameric MIS18 complex, composed of two MIS18-binding protein 1 (M18BP1^hKNL2^), four Missegregation protein 18α (MIS18α) and two MIS18β subunits (*17–22*). The hierarchical interaction of HJURP with the MIS18 complex is essential for centromere inheritance (*23, 24*). Finally, the mechanism of new CENP-A incorporation likely requires a source of energy, as it was demonstrated for other histone variants (*25*). Several chromatin remodelers have been reported to interact with kinetochore proteins (*26–31*), but their precise function at centromeres remains obscure.

**Fig. 1.**
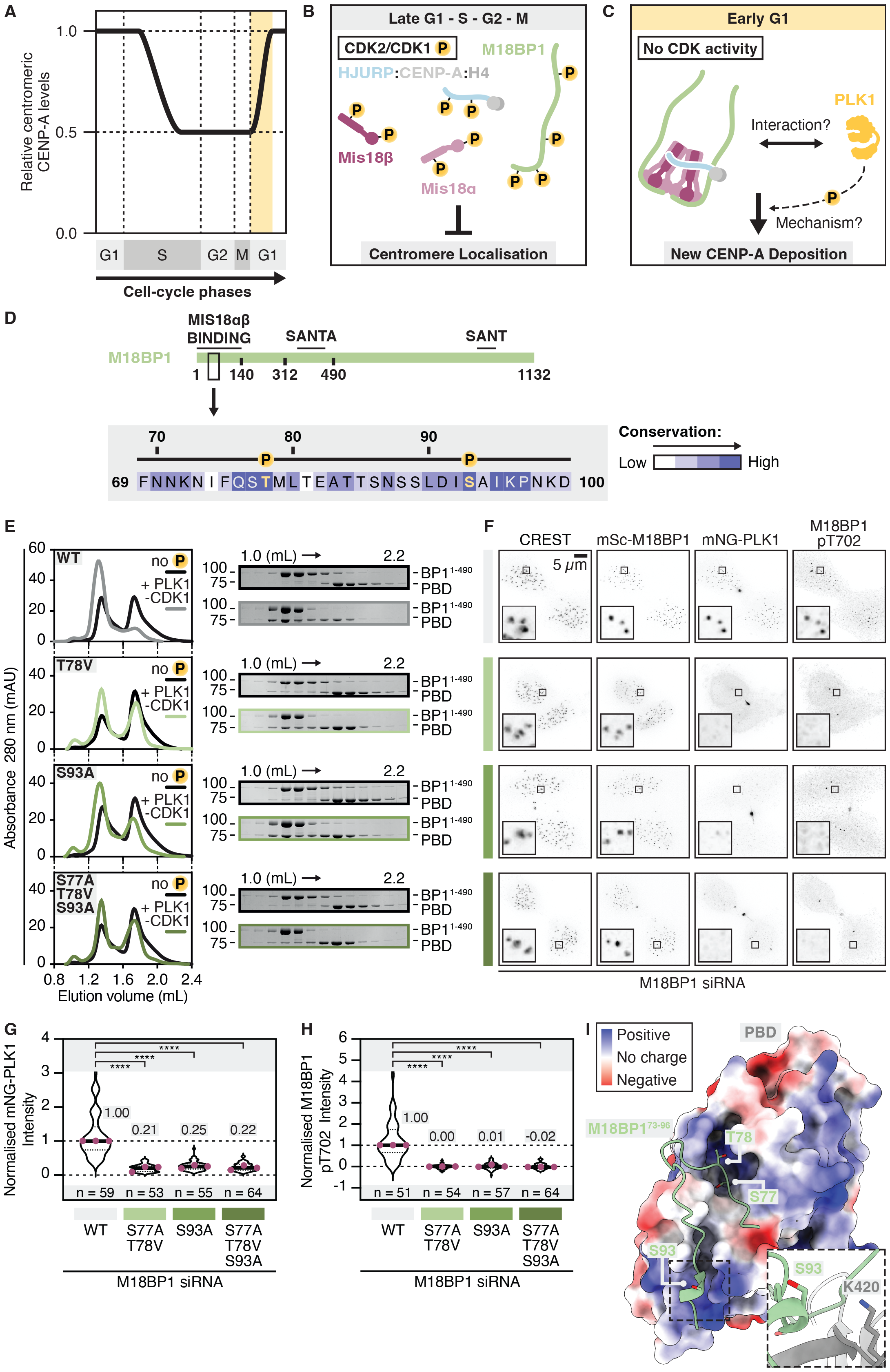
M18BP1 is the master regulator of PLK1 recruitment to centromeres in early G1. (**A**) Chromatin-bound CENP-A gets halved during DNA replication and replenished to its initial level in early G1 phase. (**B**) CDK kinases negatively regulate the assembly of the complete CENP-A deposition machinery thus preventing its centromeric localisation during late G1, S, G2 phases and mitosis. (**C**) During early G1 phase, reduced CDK activity allows the CENP-A deposition machinery to localise to centromeres. PLK1 activity is also needed to license the mechanism. (**D**) Cartoon showing the location of the PLK1 binding site on M18BP1. The inset displays the evolutionary conservation of the domain. (**E**) Analytical SEC of MBP-M18BP1^1-490^ (*wild-type* or mutants) incubated with MBP-PBD in the presence or absence of kinase activity. (**F**) Centromeric localisation of mNG-PLK1 and enrichment of M18BP1 pThr702 signal in human cells expressing mScarlet-M18BP1 (*wild-type* or mutants) in early G1 phase. (**G**) Quantification of mNG-PLK1 intensity from cells in (F). (**H**) Quantification of M18BP1 pThr702 intensity from cells in (F). Magenta dots show the median value of each experimental repeat. (**I**) AlphaFold2 model showing the docking of M18BP1’s Ser77 and Thr78 to the canonical binding pocket of PBD. Ser93 is predicted to interact with a proximal positively charged patch (highlighted in the inset).

In its outline, the CENP-A deposition process echoes the regulatory network that limits the initiation of DNA replication to only once per cell cycle. In the case of CENP-A, the process is licensed by the deactivation of CDKs at mitotic exit and early G1 phase, which otherwise act as potent inhibitors of CENP-A deposition in all the other cell cycle phases. CDKs phosphorylate a constellation of residues across the CENP-A deposition machinery, preventing its untimely assembly (**Fig. 1B-C**). Specifically, CDKs activity negatively regulate CENP-A:HJURP recruitment to centromeres (*32*), represses the interaction of HJURP with the MIS18 complex and prevents M18BP1 binding to MIS18αβ, which in turn blocks their centromere localisation (*17, 19, 33, 34*). Adding another layer of complexity to this network, PLK1 activity was reported to be essential for new CENP-A deposition (*35*). Thus, two kinases with opposing functions control the epigenetic maintenance of centromeres, but how PLK1 regulates this process remains unknown (**Fig. 1C**). In this study, we elucidate this fundamental molecular mechanism, decoding it from the multiplicity of other functions that PLK1 performs during mitotic exit (*36–39*). We show that PLK1 activity is crucial to control the conformational switch of MIS18α’s N-terminus which licenses the recruitment of HJURP to centromeres. PLK1 binds sequentially to M18BP1, MIS18α and HJURP, and its physical interaction is required to promote CENP-A deposition, thus defining it as an integral part of the deposition machinery.

### M18BP1 is the master regulator of PLK1 recruitment to centromeres during early G1 phase

PLK1 activity during mitotic exit possibly regulates the recruitment of the deposition machinery to centromeres (*35*). Importantly, co-inhibition of CDK1 and PLK1 does not rescue the requirement for PLK1 activity in this process (*35*) (**Fig. S1A-H**), indicating that the defects in CENP-A deposition are not caused by ectopic CDK1 activity upon inactivation of PLK1. To disentangle the role of PLK1 in this pathway from its other essential functions during mitotic exit, we dissected its recruitment mode to centromeres in early G1, previously reported to depend on the CENP-A deposition machinery (*35*) (**Fig. S2A-E**). The determinants of centromere recognition and maintenance of the deposition machinery lie in the first 490 residues of M18BP1 (*35*), suggesting that possible PLK1 binding sites also reside here. We used *in vitro* reconstitution to assess this hypothesis. M18BP1^1-490^ was incubated with the Polo-box-domain (PBD) of PLK1, a specific phospho-aminoacidic adaptor that promotes the docking of PLK1 to previously phosphorylated motifs (*40*). Largely substoichiometric amounts of full-length PLK1 were added to trigger phosphorylation. This strategy allowed us to use the PBD at concentrations similar to those of M18BP1^1-490^ but in the presence of limiting kinase activity to promote specific phosphorylation (*39*). Substoichiometric CDK1 was also added to the mixture, as it primes PLK1 binding to various substrates during mitosis, including the kinetochore (*41*). Complex assembly between M18BP1^1-490^ and the PBD was then examined using solid-phase assays. The results showed that PLK1 interacts with a region spanning M18BP1’s first 140 residues (**Fig. S3A**). Detailed analysis of this domain spotlighted two highly conserved residues that might participate in the interaction, Thr78 and Ser93 (**Fig. 1D**). Antibodies raised against the phosphorylated versions of these two residues showed that Thr78 is a PLK1-specific substrate, whereas Ser93 can be phosphorylated by both PLK1 and CDK1 *in vitro* (**Fig. S3B-C**). We used analytical size-exclusion chromatography (SEC) to assess the relative contributions of these kinases in triggering PLK1 binding to M18BP1. These assays demonstrated that phosphorylation by PLK1, contrary to CDK1, was largely sufficient for a stoichiometric interaction of the PBD with M18BP1^1-490^ (**Fig. S4A-C**). Mutating both Thr78 and Ser93 abolished the interaction between M18BP1^1-490^ and PBD *in vitro*, whereas the single mutations weakened it (**Fig. 1E**). We subsequently used rescue assays to study the extent of PLK1 recruitment to centromeres in human cells. In these experiments, the depletion of the endogenous M18BP1 was compensated by the expression of an inducible siRNA-resistant, fluorescently-tagged M18BP1. Interestingly, the individual mutations were sufficient to abrogate the recruitment of PLK1 to centromeres in early G1 (**Fig. 1F-G, Fig. S5A-C**). Cells expressing the M18BP1 phosphorylation mutants retained normal cytokinesis and PLK1 localisation at the spindle midbody remnant (**Fig. 1F**), but suffered from a drop of PLK1 localisation and centromere activity, as measured by visualising M18BP1’s Thr702, a known PLK1 substrate (*35*) (**Fig. 1F, H**).

SEC coupled to multi-angle light scattering (SEC-MALS) measurements demonstrated that M18BP1 and PLK1 bind in a 1:1 stoichiometry (**Fig. S6A-B**). AlphaFold2 modelling (*42*) predicts that Thr78, which is part of a canonical PBD-binding motif (S-S/T-X), docks the established phosphosite-binding pocket of PLK1’s PBD, whereas Ser93 is predicted to contact a nearby positively charged patch, thus contributing to strengthening the interaction upon phosphorylation (**Fig. 1I, S6C**). These results show that M18BP1 is the master regulator of PLK1 recruitment to centromeres during early G1 phase and describe a previously unknown binding mode to the PBD engaging two closely spaced phosphorylated residues.

### Centromeric PLK1 is required for HJURP recruitment to centromeres in early G1

Given the dramatic loss of PLK1 recruitment in early G1 displayed by the M18BP1 point mutants, we subsequently studied the effects on the localisation of the deposition machinery and CENP-A deposition. Using rescue assays, we found that M18BP1 point mutants were unable to load new CENP-A at centromeres (**Fig. 2A-B, Fig. S7A-B**). Ablation of PLK1 recruitment by the mutations had little or no effect on the centromeric localisation of M18BP1 and MIS18α, a proxy for the MIS18αβ complex (**Fig. 2A, Fig. S7C-D**). M18BP1’s binding region to MIS18αβ (residues 1-140) encompasses the PLK1-binding site. However, mutations at Thr78 and Ser93 did not affect the binding to the MIS18αβ complex in an analytical SEC experiment (**Fig. S8**). These results imply that PLK1 regulates a process downstream from the assembly of the MIS18 complex. Using M18BP1^S77A-T78V-S93A^ as the representative PLK1-loss-of-binding mutant, we examined HJURP recruitment using the same strategy as above. Crucially, expressing M18BP1^S77A-T78V-S93A^ in cells depleted of endogenous M18BP1 resulted in an almost complete loss of HJURP from centromeres (**Fig. 2C-D, Fig. S9A-C**). We conclude that, during early G1, PLK1 binds to the N-terminal region of M18BP1 and positively regulates HJURP recruitment to centromeres to promote new CENP-A deposition (**Fig. 2E**).

**Fig. 2.**
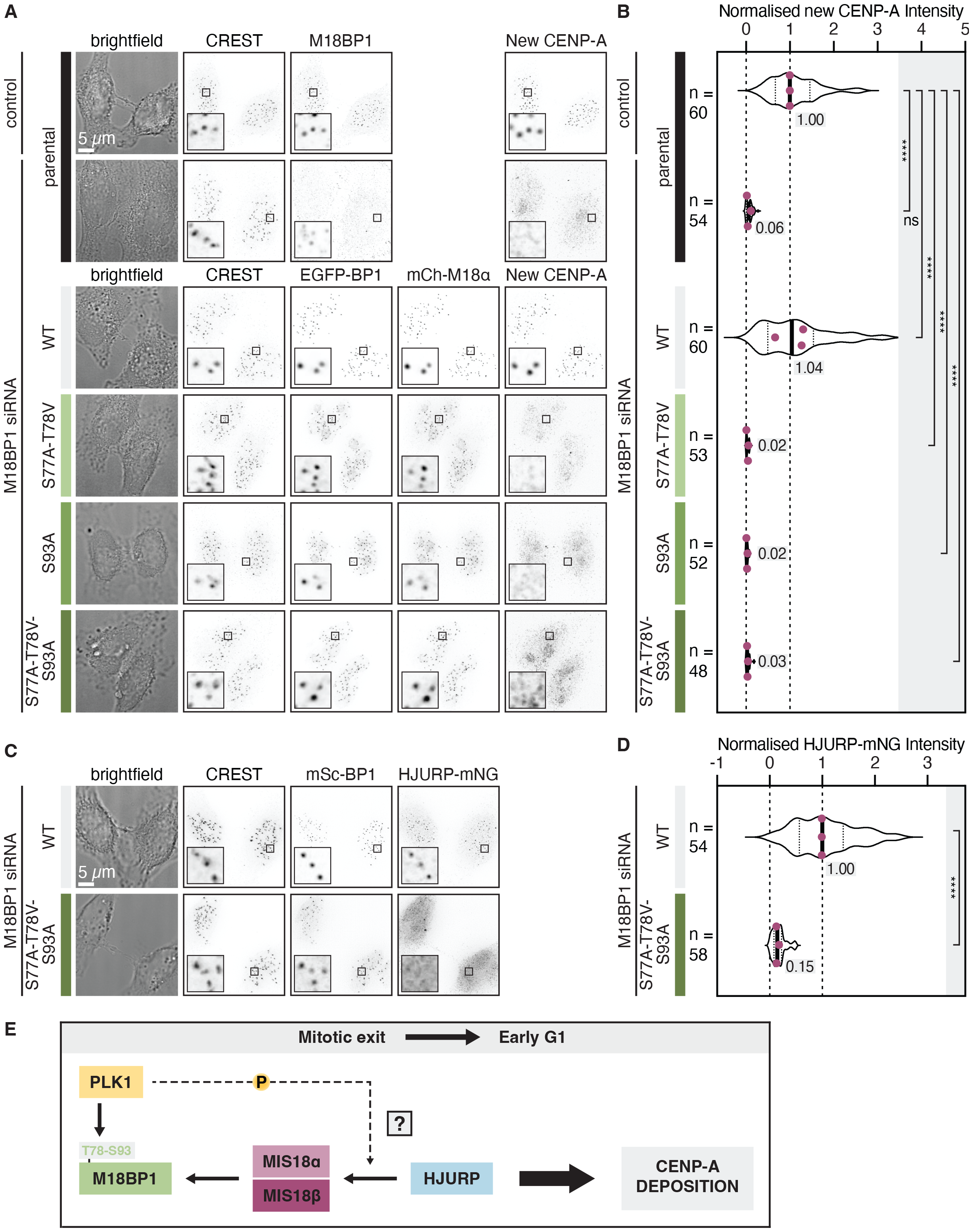
Centromeric PLK1 is required for HJURP recruitment to centromeres in early G1. (**A**) EGFP-M18BP1, mCherry-MIS18α and new CENP-A levels at centromeres in human cells expressing EGFP-M18BP1 (*wild-type* or mutants) in early G1 phase. The upper panels show the parental cells as controls for the endogenous M18BP1 depletion. (**B**) Quantification of new CENP-A levels in cells from (A). Magenta dots show the median value of each experimental repeat. (**C**) HJURP-mNG levels in human cells expressing M18BP1 (*wild-type* or S77A-T78V-S93A mutant) in early G1 phase. (**D**) Quantification of HJURP-mNG intensity in cells from (C). Magenta dots show the median value of each experimental repeat. (**E**) Summary diagram: in early G1, PLK1 binds to M18BP1’s N-terminus and regulates HJURP localisation to centromeres to allow CENP-A deposition.

### PLK1 relieves MIS18α’s N-terminal gatekeeping activity on HJURP recruitment to centromeres

The N-terminal 55 residues of MIS18α exert an inhibitory effect *in vitro* on the interaction of HJURP with the MIS18αβ complex *(23)*, possibly reflecting a conformational switch (**Fig. 3A**). We surmised that PLK1 activity could regulate this gatekeeping activity. To test this hypothesis, we devised an *in vitro* reconstitution system consisting of the MIS18αβ complex and the C-terminal region of HJURP encompassing the R2 repeat (541-C fragment), which promotes MIS18αβ binding (*23*). We performed the experiment at concentrations that allowed us to detect the possible changes in binding affinity resulting from PLK1 activity. The results revealed that PLK1 phosphorylation alone was insufficient for complex formation (**Fig. 3B** - purple trace, **Fig. S10A**). Strikingly, when the PBD was added to the system to mimic possible PLK1 binding, we detected the formation of a stable complex composed of HJURP^541-C^, MIS18αβ and PBD (**Fig. 3B** - pink trace). PBD was added at concentrations expected to saturate binding to all the possible docking sites. Complex formation was dependent on PLK1 activity. Importantly, this complex incorporated all the components above, as mixing MIS18αβ (the largest constituent) and PBD alone resulted in a smaller complex (**Fig. 3B** - compare the green and pink traces). Performing the same experiment with HJURP R1 repeat (394-540 fragment), which also binds MIS18αβ complex, yielded a similar outcome (**Fig. S10B-C**). Further dissection of this mechanism showed that PLK1 interacts with a conserved motif enclosed in MIS18α’s first 55 residues, and that binding was dependent on the phosphorylation of Ser54 of MIS18α (**Fig. 3C, Fig. S11A-B**). These results show that PLK1, contingent to phosphorylation of and binding to MIS18α, relieves a steric blockade of MIS18α’s N-terminal region on the HJURP binding site, thus promoting the formation of the HJURP:MIS18αβ complex.

**Fig. 3.**
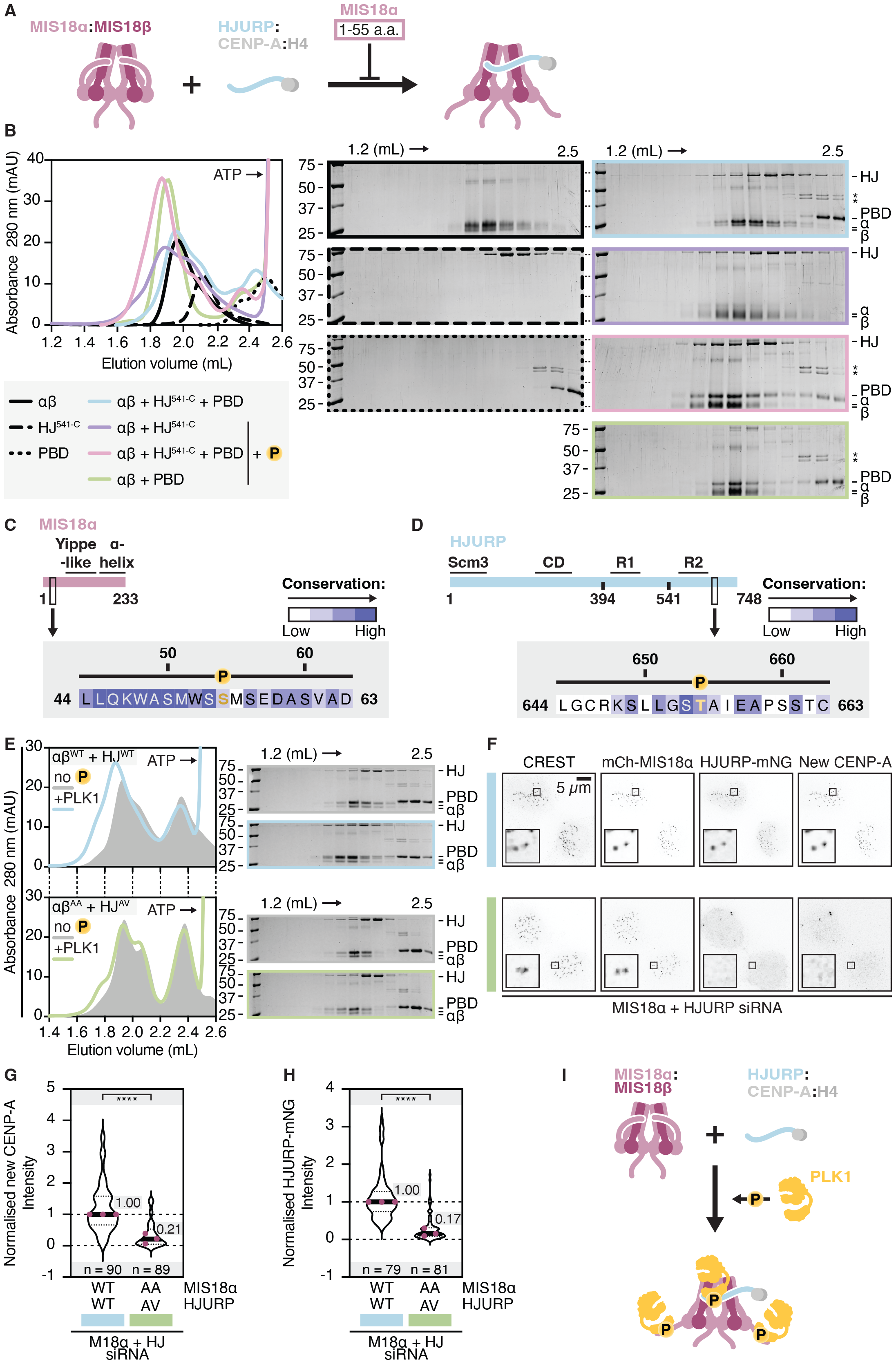
PLK1 relieves MIS18α’s N-terminal gatekeeping activity on HJURP recruitment to centromeres. (**A**) Summary diagram of previous findings: the N-terminal 55 residues of MIS18α exert a negative regulation of HJURP binding to MIS18αβ. (**B**) Analytical SEC shows that MIS18αβ and HJURP541-^C^ can bind in the presence of PBD and PLK1 phosphorylation. (**C**) Cartoon showing the location of the PLK1 binding site on MIS18α. The inset displays the evolutionary conservation of the domain. (**D**) Cartoon showing the location of the PLK1 binding site on HJURP. The inset displays the evolutionary conservation of the domain. (**E**) Analytical SEC shows that HJURP541-^C^ interaction with MIS18αβ is abrogated if the PBD is not able to bind to both of them. (**F**) mCherry-MIS18α, HJURP-mNG and new CENP-A levels in human cells co-expressing mCherry-MIS18α and HJURP-mNG (both *wild-type* or mutant) in early G1 phase. (**G**) Quantification of new CENP-A intensity from (F). (**H**) Quantification of HJURP-mNG intensity from (F). Magenta dots show the median value of each experimental repeat. (**I**) Summary diagram: PLK1 phosphorylation of and binding to MIS18α and HJURP license their interaction through MIS18α’s N-terminus conformational switch.

### PLK1 binds to both MIS18α and HJURP to license new CENP-A deposition

Our exploration of the docking of PLK1 on the HJURP:MIS18αβ complex also uncovered a conserved PBD binding domain in the HJURP C-terminal region, which required phosphorylation of Thr654 (**Fig. 3D, Fig. S12A-D**). Crucially, incubating recombinant MIS18α^S53A-S54A^β with HJURP^S653A-T654V^ abrogated the formation of the HJURP:MIS18αβ complex even in the presence of PBD and PLK1 activity (**Fig. 3E, Fig. S13A-B**). Different combinations of *wild-type* and mutant proteins resulted in complexes exhibiting intermediate affinities (**Fig. S13C-F**). To demonstrate that these interactions are essential to attain successful CENP-A deposition in a cellular context, we performed rescue assays where HJURP^S653A-T654V^ was co-expressed with MIS18α^S53A-S54A^, both fluorescently labelled, in human cells depleted of the endogenous proteins. As predicted, co-expression of the two mutants dramatically reduced the ability of the cells to perform CENP-A deposition (**Fig. 3F-G, Fig. S14A-E**) and decreased the centromeric recruitment of HJURP to levels similar to those exhibited by the M18BP1^S77A-T78V-S93A^ mutant (**Fig. 3F, H** - compare to **Fig. 2C-D**). As anticipated by the binding assays *in vitro*, the expression of individual point mutants of either MIS18α or HJURP only mildly affected CENP-A deposition (**Fig. S15A-L**). We conclude that simultaneous binding of PLK1 to MIS18α and HJURP is needed to counteract the gatekeeping activity of MIS18α N-terminus and license successful CENP-A deposition (**Fig. 3I**).

### The role of PLK1 in the epigenetic maintenance of centromeres revolves around the MIS18α conformational switch

PLK1 licensing activity lies in its ability to regulate the MIS18α N-terminal conformation (**Fig. 3**). If this were the primary role of PLK1 in the pathway of CENP-A deposition, then removing MIS18α’s inhibitory domain ought to be able to sustain CENP-A deposition in the absence of PLK1 activity at centromeres. To verify this hypothesis, we designed a rescue assay in human cells where MIS18α^56-C^ was co-expressed with M18BP1^S77A-T78V-S93A^ to abrogate PLK1 recruitment to centromeres (**Fig. S16A**). The endogenous proteins were knocked down by siRNA-mediated depletion. As shown above, expression of M18BP1^S77A-T78V-S93A^ abrogated CENP-A deposition (**Fig. 4A-B, Fig. S16B-F**). Critically, cells expressing MIS18α^56-C^ alongside M18BP1^S77A-T78V-S93A^ incorporated new CENP-A to substantial levels (58% efficiency in comparison to the *wild-type* control - **Fig. 4A-B**). Altogether, these results demonstrate that the role of PLK1 in new CENP-A deposition impinges on the regulation of the MIS18α conformational switch.

**Fig. 4.**
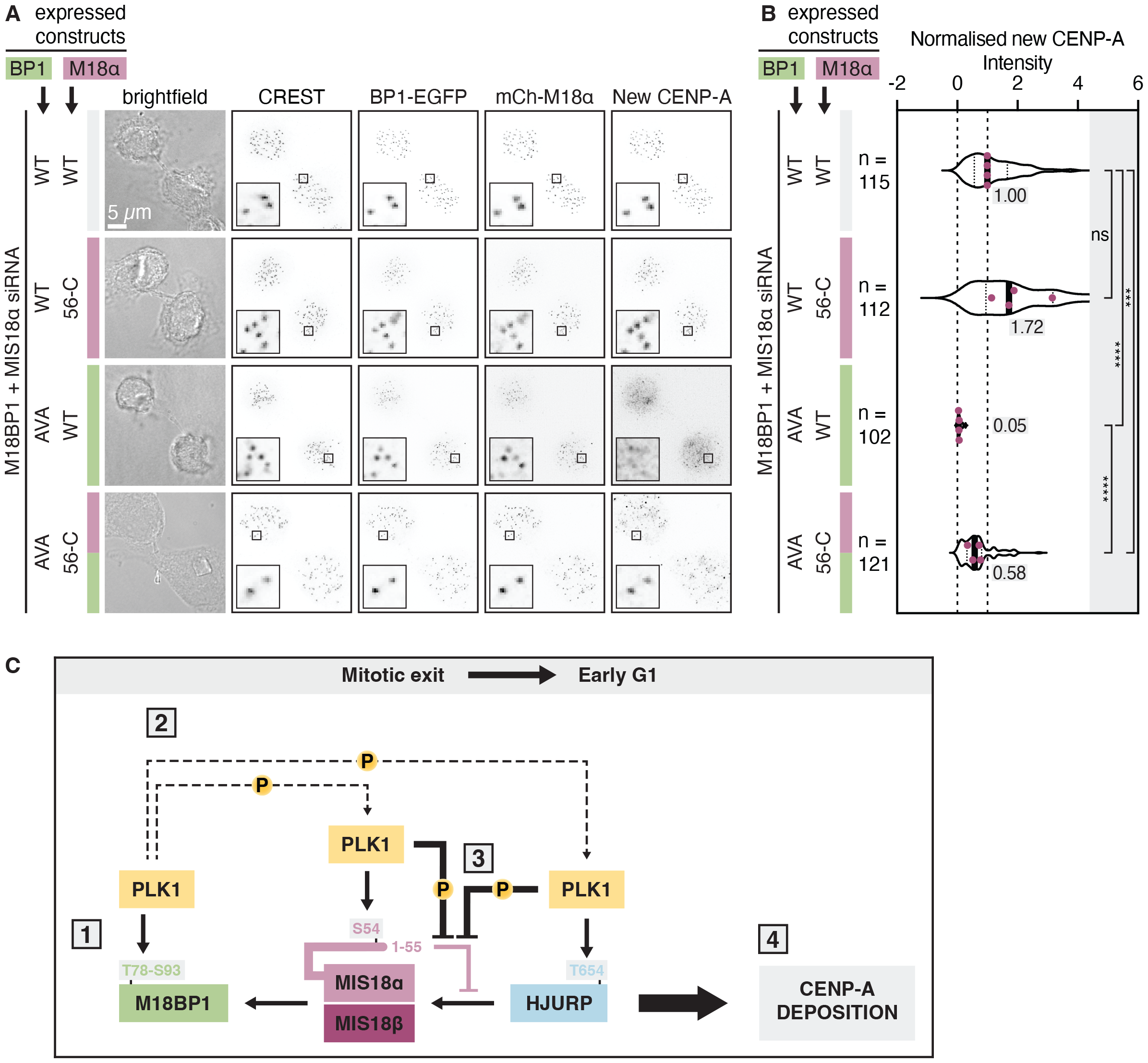
The role of PLK1 in the epigenetic maintenance of centromeres revolves around MIS18α’s conformational switch. (**A**) EGFP-M18BP1, mCherry-MIS18α and new CENP-A levels in human cells co-expressing EGFP-M18BP1 and mCherry-MIS18α (*wild-type* or mutants, in different combinations) in early G1 cells. (**B**) Quantification of new CENP-A intensity from (A). Magenta dots show the median value of each experimental repeat. (**C**) Diagram showing the final model of this study.

### PLK1 binding to the CENP-A deposition machinery is hierarchical

Our work shows that PLK1 regulates CENP-A deposition through multiple interactions with the components of the deposition machinery. Since mutating the PLK1-docking site on M18BP1 was sufficient to prevent HJURP recruitment to centromeres, our data suggests a possible binding hierarchy promoting sequential recruitment of the various components of the deposition machinery in cells (*15, 18*). Then, we asked whether grafting this domain onto HJURP could sustain its centromere localisation and CENP-A deposition. A minimal PLK1-binding fragment of M18BP1 encompassing the 56-98 residues bound to PLK1 in analytical SEC experiments (**Fig. S17A-B**).

We used this fragment to replace the PBD-binding motif of HJURP (**Fig. S18A** - this chimeric construct is henceforth called “HJURP^graft^”), and co-expressed HJURP^graft^ with the M18BP1 ^S77A-T78V-S93A^ mutant in human cells, all in the background of the endogenous proteins depletion (**Fig. S18B**). The HJURP^graft^ construct did not localise to centromeres and did not allow new CENP-A deposition in the presence of the M18BP1^S77A-T78V-S93A^ mutant (**Fig. S18C-G**). Consistent with these results, preventing the binding of PLK1 to another component of the deposition machinery downstream of M18BP1 recruitment, MIS18α (S53A-S54A mutant), barely affected PLK1 enrichment to centromeres (**Fig. S19A-G**). In addition, preventing HJURP from interacting with PLK1 did not significantly alter its localisation to centromeres (**Fig. S15I, K**). Therefore, PLK1 licensing of new CENP-A deposition must start with its binding to M18BP1.

## Discussion

Our work solves the long-standing question of the role of PLK1 kinase in licensing new CENP-A deposition (*35*). In early G1, PLK1 binds to M18BP1 on a highly conserved binding site generated by the phosphorylation of two nearby residues (Thr78 and Ser93) (**Fig. 4C**, Step 1), both of which have to be phosphorylated for successful recruitment of PLK1 to centromeres. This doubly phosphorylated site interacts with a single PBD, a previously undescribed binding mode that expands our collective knowledge of how PLK1 docks on its targets (*40, 43*). M18BP1’s Ser93 is a substrate of both PLK1 and CDK1 *in vitro*. Its sequence resembles non-canonical CDK1 sites previously identified in other mitotic substrates (*44–46*) but also conforms to a classical acidophilic PLK1 substrate. This promiscuous regulation hints at possible priming of the binding during mitosis, in parallel with other mitotic PLK1-binding proteins (*41*), implying that the phosphorylated site needs to survive dephosphorylation by PP2A-B55 during mitotic exit (*47*). The antibodies against the phosphorylated version of M18BP1’s Thr78 and Ser93 generated in this work yielded poor results when used *in vivo* to assess the dynamics of phosphorylation in the transition from mitotic exit to early G1 phase (DC and AM, unpublished results).

To overcome pleiotropic effects from using PLK1 inhibitors, we focused on specific phosphorylation sites and separation-of-function mutants. This brought to light that centromere targeting of M18BP1 and the MIS18αβ complex is minimally sensitive to PLK1 activity, but uncovered a complex cascade of events that ultimately activate CENP-A discharge by the deposition machinery. We hypothesise that once bound to M18BP1, PLK1 additionally primes the simultaneous binding of additional PLK1 molecules to two downstream components of the CENP-A deposition machinery (**Fig. 4C**, Step 2). The interactions require the phosphorylation of Ser54 on MIS18α and Thr654 on HJURP. PLK1 physical binding is essential for increasing the affinity between HJURP and MIS18αβ, a finding that explains how MIS18α’s N-terminal α-helix gatekeeping activity is relieved (*23*); PLK1 interaction stabilises a conformational switch which, in turn, enhances the affinity between HJURP and MIS18αβ (**Fig. 4C**, Step 3). All the PLK1-docking sites mapped in this study are located in proximity to the regions that the deposition machinery components use to interact with each other (*17, 19, 23, 24*) (**Fig. S20**). Thus, CENP-A deposition appears to require full assembly of this complex (**Fig. 4C**, Step 4). By showing that PLK1 is a tightly docked component, our data imply a structural role of PLK1 in the assembly of the complex. The regulation of the MIS18α’s N-terminal region is pivotal for HJURP localisation to centromeres, as it is corroborated by the MIS18α^56-C^ mutant being able to execute CENP-A deposition in the absence of PLK1 at centromeres.

The recruitment of PLK1 onto the CENP-A deposition machinery appears to be sequential, and it depends on binding first to M18BP1, which acts as its master regulator. The hierarchy of binding is reflected by the fact that M18BP1’s PLK1-binding site cannot be grafted onto another downstream protein without losing CENP-A deposition activity. This order of binding is consistent with the sequential centromere localisation of the CENP-A deposition machinery in human cells during the transition from mitotic exit to early G1 (*15, 18*). The conservation of the PLK1 sites implies that this mechanism could also work in other metazoans. The importance of M18BP1:PLK1 binding correlates to the mild CENP-A deposition phenotypes exhibited by the single expression of MIS18α^S53A-S54A^ and HJURP^S653A-T654V^ mutants. The dramatic loss of CENP-A incorporation exhibited by the M18BP1^S77A-T78V-S93A^ mutant was indeed phenocopied only when both MIS18α^S53A-S54A^ and HURP^S653A-T654V^ mutants were co-expressed. Our analysis does not exclude that PLK1, in addition to controlling the incorporation of HJURP in the MIS18 complex, may regulate further downstream steps in the mechanism of new CENP-A incorporation.

Our study sheds new insights into how cells replenish the centromere epigenetic marker at every cell cycle. By employing two kinases (CDK1 and PLK1) with opposite functions, the system ensures that the CENP-A deposition machinery does not assemble untimely. This safety mechanism needs indeed to satisfy two conditions to allow the deposition reaction to happen; i) CDK1 sites must be dephosphorylated (*17, 32–34*) and ii) PLK1 must phosphorylate and bind to M18BP1, MIS18α and HJURP (*35*). This elegant solution restricts the deposition reaction to the early G1 phase. Our findings are consistent with and complementary to the work of *Parashara, Medina-Pritchard, Abad* and colleagues.

## Acknowledgements

We thank all the members of the Musacchio group for the helpful scientific discussion. R.G. Schönenbrücher and P. Geue (MPI Dortmund, Biophysics Facility) for support with SEC-MALS. P. Keller (MPI Dresden, Antibody Facility) for helping with antibody generation. C. Körner, A. Dammers, L. Schulze and L. Oberste-Lehn for assisting with reagent preparation. N. Schmidt for support with image analysis. S. Maffini and J. Schweighofer for critical reading of the manuscript. A.A. Jeyaprakash, P. Parashara, B. Medina-Pritchard and M.A. Abad for sharing unpublished results and discussion during the development of this study.

## Funding

Max Planck Society (AM)

European Research Council Synergy Grant 951430 BIOMECANET (AM)

Marie-Curie Training Network DivIDE project number 675737 (AM)

DGF’s Collaborative Research Centre 1430 “Molecular Mechanisms of Cell State Transitions” (AM)

CANTAR network under the Netzwerke-NRW programme (AM)

EMBO Long-Term Fellowship ALFT439-2019 (DC)

## Author contributions

Conceptualization: AM, DC

Methodology: DC, AEV

Investigation: DC, AEV, IRV, DP, VC, MP

Visualization: DC

Funding acquisition: AM

Project administration: AM, DC

Supervision: AM

Writing – original draft: DC

Writing – review & editing: AM, DC, AEV, IV

## Data and materials availability

All data are available in the main text or the supplementary materials.

## Supplementary Materials

Materials and Methods

Supplementary Text

Figs. S1 to S20

Tables S1 to S5

References (*48–58*)

## References

1. A. Musacchio, A. Desai, A Molecular View of Kinetochore Assembly and Function. Biology. 6 (2017), doi:10.3390/biology6010005.

2. E. Zasadzińska, D. R. Foltz, Orchestrating the Specific Assembly of Centromeric Nucleosomes. Prog. Mol. Subcell. Biol. 56, 165–192 (2017).

3. T. Fukagawa, W. C. Earnshaw, The centromere: chromatin foundation for the kinetochore machinery. Dev. Cell. 30, 496–508 (2014).

4. J. Ohzeki, V. Larionov, W. C. Earnshaw, H. Masumoto, De novo formation and epigenetic maintenance of centromere chromatin. Curr. Opin. Cell Biol. 58, 15–25 (2019).

5. K. L. McKinley, I. M. Cheeseman, The molecular basis for centromere identity and function. Nat. Rev. Mol. Cell Biol. 17, 16–29 (2016).

6. P. B. Talbert, S. Henikoff, What makes a centromere? Exp. Cell Res. 389, 111895 (2020).

7. O. J. Marshall, A. C. Chueh, L. H. Wong, K. H. A. Choo, Neocentromeres: new insights into centromere structure, disease development, and karyotype evolution. Am. J. Hum. Genet. 82, 261–282 (2008).

8. K. F. Sullivan, M. Hechenberger, K. Masri, Human CENP-A contains a histone H3 related histone fold domain that is required for targeting to the centromere. J. Cell Biol. 127, 581–592 (1994).

9. K. Yoda, S. Ando, S. Morishita, K. Houmura, K. Hashimoto, K. Takeyasu, T. Okazaki, Human centromere protein A (CENP-A) can replace histone H3 in nucleosome reconstitution in vitro. Proceedings of the National Academy of Sciences. 97, 7266–7271 (2000).

10. D. Fachinetti, H. D. Folco, Y. Nechemia-Arbely, L. P. Valente, K. Nguyen, A. J. Wong, Q. Zhu, A. J. Holland, A. Desai, L. E. T. Jansen, D. W. Cleveland, A two-step mechanism for epigenetic specification of centromere identity and function. Nat. Cell Biol. 15, 1056–1066 (2013).

11. E. M. Dunleavy, G. Almouzni, G. H. Karpen, H3.3 is deposited at centromeres in S phase as a placeholder for newly assembled CENP-A in G1 phase. Nucleus. 2, 146–157 (2011).

12. D. L. Bodor, L. P. Valente, J. F. Mata, B. E. Black, L. E. T. Jansen, Assembly in G1 phase and long-term stability are unique intrinsic features of CENP-A nucleosomes. Mol. Biol. Cell. 24, 923–932 (2013).

13. L. E. T. Jansen, B. E. Black, D. R. Foltz, D. W. Cleveland, Propagation of centromeric chromatin requires exit from mitosis. J. Cell Biol. 176, 795–805 (2007).

14. M. Schuh, C. F. Lehner, S. Heidmann, Incorporation of Drosophila CID/CENP-A and CENP-C into centromeres during early embryonic anaphase. Curr. Biol. 17, 237–243 (2007).

15. E. M. Dunleavy, D. Roche, H. Tagami, N. Lacoste, D. Ray-Gallet, Y. Nakamura, Y. Daigo, Y. Nakatani, G. Almouzni-Pettinotti, HJURP is a cell-cycle-dependent maintenance and deposition factor of CENP-A at centromeres. Cell. 137, 485–497 (2009).

16. D. R. Foltz, L. E. T. Jansen, A. O. Bailey, J. R. Yates 3rd, E. A. Bassett, S. Wood, B. E. Black, D. W. Cleveland, Centromere-specific assembly of CENP-a nucleosomes is mediated by HJURP. Cell. 137, 472–484 (2009).

17. D. Pan, K. Klare, A. Petrovic, A. Take, K. Walstein, P. Singh, A. Rondelet, A. W. Bird, A. Musacchio, CDK-regulated dimerization of M18BP1 on a Mis18 hexamer is necessary for CENP-A loading. Elife. 6 (2017), doi:10.7554/eLife.23352.

18. Y. Fujita, T. Hayashi, T. Kiyomitsu, Y. Toyoda, A. Kokubu, C. Obuse, M. Yanagida, Priming of Centromere for CENP-A Recruitment by Human hMis18α, hMis18β, and M18BP1. Dev. Cell. 12, 17–30 (2007).

19. F. Spiller, B. Medina-Pritchard, M. A. Abad, M. A. Wear, O. Molina, W. C. Earnshaw, A. Jeyaprakash, Molecular basis for Cdk1-regulated timing of Mis18 complex assembly and CENP-A deposition. EMBO Rep. 18, 894–905 (2017).

20. R. Thamkachy, B. Medina-Pritchard, S. H. Park, C. G. Chiodi, J. Zou, M. de la Torre-Barranco, K. Shimanaka, M. A. Abad, C. G. Páramo, R. Feederle, E. Ruksenaite, P. Heun, O. R. Davies, J. Rappsilber, D. Schneidman-Duhovny, U.-S. Cho, A. Arockia Jeyaprakash, Structural Basis for Mis18 Complex Assembly: Implications for Centromere Maintenance. bioRxiv (2023), p. 2021.11.08.466737.

21. I. K. Nardi, E. Zasadzińska, M. E. Stellfox, C. M. Knippler, D. R. Foltz, Licensing of Centromeric Chromatin Assembly through the Mis18α-Mis18β Heterotetramer. Mol. Cell. 61, 774–787 (2016).

22. M. E. Stellfox, I. K. Nardi, C. M. Knippler, D. R. Foltz, Differential Binding Partners of the Mis18α/β YIPPEE Domains Regulate Mis18 Complex Recruitment to Centromeres. Cell Rep. 15, 2127–2135 (2016).

23. D. Pan, K. Walstein, A. Take, D. Bier, N. Kaiser, A. Musacchio, Mechanism of centromere recruitment of the CENP-A chaperone HJURP and its implications for centromere licensing. Nat. Commun. 10, 4046 (2019).

24. J. Wang, X. Liu, Z. Dou, L. Chen, H. Jiang, C. Fu, Mitotic regulator Mis18β interacts with and specifies the centromeric assembly of molecular chaperone HJURP. Journal of Biological (2014) (available at http://www.jbc.org/content/early/2014/02/11/jbc.M113.529958.short).

25. G. Mizuguchi, X. Shen, J. Landry, W.-H. Wu, S. Sen, C. Wu, ATP-driven exchange of histone H2AZ variant catalyzed by SWR1 chromatin remodeling complex. Science. 303, 343–348 (2004).

26. D. R. Foltz, L. E. T. Jansen, B. E. Black, A. O. Bailey, J. R. Yates 3rd, D. W. Cleveland, The human CENP-A centromeric nucleosome-associated complex. Nat. Cell Biol. 8, 458–469 (2006).

27. J. Walfridsson, P. Bjerling, M. Thalen, E.-J. Yoo, S. D. Park, K. Ekwall, The CHD remodeling factor Hrp1 stimulates CENP-A loading to centromeres. Nucleic Acids Res. 33, 2868–2879 (2005).

28. H. Izuta, M. Ikeno, N. Suzuki, T. Tomonaga, N. Nozaki, C. Obuse, Y. Kisu, N. Goshima, F. Nomura, N. Nomura, K. Yoda, Comprehensive analysis of the ICEN (Interphase Centromere Complex) components enriched in the CENP-A chromatin of human cells. Genes Cells. 11, 673–684 (2006).

29. M. Perpelescu, N. Nozaki, C. Obuse, H. Yang, K. Yoda, Active establishment of centromeric CENP-A chromatin by RSF complex. J. Cell Biol. 185, 397–407 (2009).

30. M. C. Barnhart-Dailey, P. Trivedi, P. T. Stukenberg, D. R. Foltz, HJURP interaction with the condensin II complex during G1 promotes CENP-A deposition. Mol. Biol. Cell. 28, 54–64 (2017).

31. M. Okada, K. Okawa, T. Isobe, T. Fukagawa, CENP-H–containing Complex Facilitates Centromere Deposition of CENP-A in Cooperation with FACT and CHD1. MBoC. 20, 3986–3995 (2009).

32. S. Müller, R. Montes de Oca, N. Lacoste, F. Dingli, D. Loew, G. Almouzni, Phosphorylation and DNA Binding of HJURP Determine Its Centromeric Recruitment and Function in CenH3CENP-A Loading. Cell Rep. 8, 190–203 (2014).

33. M. C. C. Silva, D. L. Bodor, M. E. Stellfox, N. M. C. Martins, H. Hochegger, D. R. Foltz, L. E. T. Jansen, Cdk activity couples epigenetic centromere inheritance to cell cycle progression. Dev. Cell. 22, 52–63 (2012).

34. A. Stankovic, L. Y. Guo, J. F. Mata, D. L. Bodor, X.-J. Cao, A. O. Bailey, J. Shabanowitz,D. F. Hunt, B. A. Garcia, B. E. Black, L. E. T. Jansen, A Dual Inhibitory Mechanism Sufficient to Maintain Cell-Cycle-Restricted CENP-A Assembly. Mol. Cell. 65, 231–246 (2017).

35. K. L. McKinley, I. M. Cheeseman, Polo-like kinase 1 licenses CENP-A deposition at centromeres. Cell. 158, 397–411 (2014).

36. M. Petronczki, P. Lénárt, J.-M. Peters, Polo on the Rise—from Mitotic Entry to Cytokinesis with Plk1. Dev. Cell. 14, 646–659 (2008).

37. V. Archambault, D. M. Glover, Polo-like kinases: conservation and divergence in their functions and regulation. Nat. Rev. Mol. Cell Biol. 10, 265–275 (2009).

38. G. Combes, I. Alharbi, L. G. Braga, S. Elowe, Playing polo during mitosis: PLK1 takes the lead. Oncogene. 36, 4819–4827 (2017).

39. S. Zitouni, C. Nabais, S. C. Jana, A. Guerrero, M. Bettencourt-Dias, Polo-like kinases: structural variations lead to multiple functions. Nat. Rev. Mol. Cell Biol. 15, 433–452 (2014).

40. A. E. H. Elia, P. Rellos, L. F. Haire, J. W. Chao, F. J. Ivins, K. Hoepker, D. Mohammad, L. C. Cantley, S. J. Smerdon, M. B. Yaffe, The molecular basis for phosphodependent substrate targeting and regulation of Plks by the Polo-box domain. Cell. 115, 83–95 (2003).

41. P. Singh, M. E. Pesenti, S. Maffini, S. Carmignani, M. Hedtfeld, A. Petrovic, A. Srinivasamani, T. Bange, A. Musacchio, BUB1 and CENP-U, Primed by CDK1, Are the Main PLK1 Kinetochore Receptors in Mitosis. Mol. Cell. 81, 67–87.e9 (2021).

42. J. Jumper, R. Evans, A. Pritzel, T. Green, M. Figurnov, O. Ronneberger, K. Tunyasuvunakool, R. Bates, A. Žídek, A. Potapenko, A. Bridgland, C. Meyer, S. A. A. Kohl, A. J. Ballard, A. Cowie, B. Romera-Paredes, S. Nikolov, R. Jain, J. Adler, T. Back, S. Petersen, D. Reiman, E. Clancy, M. Zielinski, M. Steinegger, M. Pacholska, T. Berghammer, S. Bodenstein, D. Silver, O. Vinyals, A. W. Senior, K. Kavukcuoglu, P. Kohli, D. Hassabis, Highly accurate protein structure prediction with AlphaFold. Nature. 596, 583–589 (2021).

43. A. E. H. Elia, L. C. Cantley, M. B. Yaffe, Proteomic screen finds pSer/pThr-binding domain localizing Plk1 to mitotic substrates. Science. 299, 1228–1231 (2003).

44. P. J. Huis In ‘t Veld, S. Jeganathan, A. Petrovic, P. Singh, J. John, V. Krenn, F. Weissmann, T. Bange, A. Musacchio, Molecular basis of outer kinetochore assembly on CENP-T. Elife. 5 (2016), doi:10.7554/eLife.21007.

45. K. Suzuki, K. Sako, K. Akiyama, M. Isoda, C. Senoo, N. Nakajo, N. Sagata, Identification of non-Ser/Thr-Pro consensus motifs for Cdk1 and their roles in mitotic regulation of C2H2 zinc finger proteins and Ect2. Sci. Rep. 5, 1–9 (2015).

46. A. Al-Rawi, E. Kaye, S. Korolchuk, J. A. Endicott, T. Ly, Cyclin A and Cks1 promote kinase consensus switching to non-proline-directed CDK1 phosphorylation. Cell Rep. 42, 112139 (2023).

47. J. Holder, E. Poser, F. A. Barr, Getting out of mitosis: spatial and temporal control of mitotic exit and cytokinesis by PP1 and PP2A. FEBS Lett. 593, 2908–2924 (2019).

48. P. J. Huis In ‘t Veld, S. Wohlgemuth, C. Koerner, F. Müller, P. Janning, A. Musacchio, Reconstitution and use of highly active human CDK1:Cyclin-B:CKS1 complexes. Protein Sci. 31, 528–537 (2022).

49. R. Evans, M. O’Neill, A. Pritzel, N. Antropova, A. Senior, T. Green, A. Žídek, R. Bates, S. Blackwell, J. Yim, O. Ronneberger, S. Bodenstein, M. Zielinski, A. Bridgland, A. Potapenko, A. Cowie, K. Tunyasuvunakool, R. Jain, E. Clancy, P. Kohli, J. Jumper, D. Hassabis, Protein complex prediction with AlphaFold-Multimer. bioRxiv (2022), p. 2021.10.04.463034.

50. UniProt Consortium, UniProt: the Universal Protein Knowledgebase in 2023. Nucleic Acids Res. 51, D523–D531 (2023).

51. A. Tighe, V. L. Johnson, S. S. Taylor, Truncating APC mutations have dominant effects on proliferation, spindle checkpoint control, survival and chromosome stability. J. Cell Sci. 117, 6339–6353 (2004).

52. A. Tighe, O. Staples, S. Taylor, Mps1 kinase activity restrains anaphase during an unperturbed mitosis and targets Mad2 to kinetochores. J. Cell Biol. 181, 893–901 (2008).

53. J. Schindelin, I. Arganda-Carreras, E. Frise, V. Kaynig, M. Longair, T. Pietzsch, S. Preibisch, C. Rueden, S. Saalfeld, B. Schmid, J.-Y. Tinevez, D. J. White, V. Hartenstein, K. Eliceiri, P. Tomancak, A. Cardona, Fiji: an open-source platform for biological-image analysis. Nat. Methods. 9, 676 (2012).

54. D. L. Bodor, M. G. Rodríguez, N. Moreno, L. E. T. Jansen, Analysis of protein turnover by quantitative SNAP-based pulse-chase imaging. Curr. Protoc. Cell Biol. Chapter 8, Unit8.8 (2012).

55. S. J. Lord, K. B. Velle, R. D. Mullins, L. K. Fritz-Laylin, SuperPlots: Communicating reproducibility and variability in cell biology. J. Cell Biol. 219 (2020), doi:10.1083/jcb.202001064.

56. M. Goujon, H. McWilliam, W. Li, F. Valentin, S. Squizzato, J. Paern, R. Lopez, A new bioinformatics analysis tools framework at EMBL-EBI. Nucleic Acids Res. 38, W695–W699 (2010).

57. F. Sievers, A. Wilm, D. Dineen, T. J. Gibson, K. Karplus, W. Li, R. Lopez, H. McWilliam, M. Remmert, J. S. O. Ding, J. D. Thompson, D. G. Higgins, Fast, scalable generation of high-quality protein multiple sequence alignments using Clustal Omega. Mol. Syst. Biol. 7, 1–6 (2011).

58. A. M. Waterhouse, J. B. Procter, D. M. A. Martin, M. Clamp, G. J. Barton, Jalview Version 2--a multiple sequence alignment editor and analysis workbench. Bioinformatics. 25, 1189–1191 (2009).

