## Supplementary Information for "Role of PLK1 in the epigenetic maintenance of centromeres"

### Materials and Methods

Protein purification

All recombinant M18BP1, MIS18α:MIS18β and HJURP proteins used in this study were purified following the same 3-step methodology, a modified protocol from previous strategies (*17*, *23*). For a complete list of the plasmids used for protein expression see **Table S1**. The purification of M18BP1 fragments was performed at pH 7.5, whereas a pH of 8 and 6.9 was used for MIS18α:MIS18β and HJURP proteins, respectively. Plasmids were transformed in *E. coli* BL21-CodonPlus(DE3)-RIL cells (Agilent Technologies, #230240), and cells were grown in Terrific Broth at 37°C. For all purifications, cells were grown in 6 L of media. When the culture reached an $OD_{600}$ ranging from 0.6-0.8. 0.2 mM IPTG was added to the culture and cells were grown at 20°C for 16 hours. Cells were harvested and resuspended in lysis buffer (20 mM HEPES, 300 mM NaCl, 25 mM imidazole, 1 mM TCEP) supplemented with protease inhibitor cocktail (Serva) and DNase (Roche), lysed using a fluidiser and cleared by centrifugation at 30,000 rpm for 45 minutes at 4°C. The cleared lysate was loaded into a 5 mL HisTrap FF column (GE Healthcare) or a 5 mL GSTrap FF column (GE Healthcare), washed with 30 mL of lysis buffer, and eluted using lysis buffer containing 300 mM imidazole. The protein fractions were pooled. For the purification of the MIS18α:MIS18β complex, affinity tags were removed by digestion with TEV protease (produced in-house) overnight at 4°C. The proteins were loaded, according to their isoelectric point, into either a 5 mL HiTrap Heparin HP column (Cytiva) or a 6 mL RESOURCE Q column (Cytiva) for ion exchange chromatography. Before loading, the buffer was either diluted or exchanged by dialysis to lower the salt concentration to 20-50 mM. After loading, the proteins were eluted using a linear gradient of 20-1000 mM NaCl using the IEX elution buffer (20 mM HEPES, 1 M NaCl, 2.5% glycerol, 1 mM TCEP). The fractions containing the protein of interest were pooled, concentrated and loaded, according to their molecular weight, into either a Superdex200 16/60 (Ge Healthcare) or a HiLoad 16/600 Superose6 (Cytiva) column. The size exclusion chromatography used a buffer containing 20 mM HEPES, 300 mM NaCl, 2.5% glycerol and 1 mM TCEP. The fractions containing the protein of interest were concentrated, flash-frozen in liquid nitrogen and stored at -80°C.

MBP-TEV-BORA$^{1\text{-}224}$-8His was expressed in *E. coli* cells as above. After harvesting, cells were resuspended in lysis buffer (20 mM Tris pH8, 500 mM NaCl, 0.1% Triton X-100, 1 mM TCEP) supplemented with protease inhibitor cocktail (Serva) and DNase (Roche) and lysed by sonication (60 pulses at 1 second on, 3 seconds off, 50% amplitude). Lysates were cleared by 30 minutes of centrifugation at 25,000 g. The cleared lysate was bound for 2 hours to 3 mL of equilibrated Ni-NTA agarose slurry (QIAGEN), washed in a 10 mL gravity flow column (ThermoFisher) with buffer containing 20 mM Tris pH8, 500 mM NaCl, 20 mM Imidazole and 1 mM TCEP, and eluted using the same buffer containing 300 mM imidazole. The eluted protein was concentrated and loaded into a Superdex200 16/60 (Ge Healthcare) equilibrated with lysis buffer. The fractions containing the protein of interest were concentrated, flash-frozen in liquid nitrogen and stored at -80°C.

His-PLK1, MBP-PBD, PBD and CDK1:CyclinB1:CKS1 kinases were purified as before (*41*, *48*).

His-AURORA A was expressed in *E. coli* cells as above (cells were grown in 1 L of TB media). After harvesting, cells were resuspended in lysis buffer (50 mM HEPES pH7.5, 300 mM NaCl, 5% Glycerol, 1 mM TCEP) supplemented with 1 mM PMSF and lysed by sonication (50 pulses at 1 second on, 2 seconds off). Lysates were cleared by 30 minutes of centrifugation at 75,000 g. The cleared lysate was loaded into a 5 mL HisTrap FF column (Cytiva), washed with 30 mL of lysis buffer supplemented with 1 mM ATP, and eluted using lysis buffer containing 300 mM imidazole. The eluted protein was concentrated and loaded into a Superdex200 16/60 (Cytiva) equilibrated with lysis buffer. The fractions containing the protein of interest were concentrated, flash-frozen in liquid nitrogen and stored at -80°C.

Protein phosphorylation

For all phosphorylation reactions, kinases were used in a 1:30 molar ratio with the substrates. All the proteins of the experiment were mixed in the experiment buffer (see "solid-phase assays" and "analytical size exclusion chromatography" sections) supplemented with 2 mM ATP and 10 mM $MgCl_2$. For phosphorylation reactions using PLK1 kinase, the reaction buffer also incorporated AURORA A kinase and $BORA^{1-224}$ in a 1:1 and 1:2 molar ratio with PLK1, respectively, to allow full activation of PLK1. When PLK1 was used in stoichiometric amounts with the other reaction substrates, the activation mix (AURORA A and $BORA^{1-224}$) was not used. All phosphorylation reactions were carried out overnight in a Thermomix (Eppendorf) at 10°C at 450 rpm.

Solid-phase assays

For the solid-phase assays, proteins were diluted to either 3 or 5 μM and mixed in a buffer containing 20 mM HEPES (pH according to the "protein purification" section), 350 mM NaCl, 2.5% glycerol and 1 mM TCEP, with or without the kinase mix. After the kinase reaction was completed, proteins were mixed with 20 μL of equilibrated amylose (NEB) or glutathione (manufacturer) resin. One-third of the mixture was taken as input fraction and the rest was incubated in a Thermomix (NEB) at 4°C for 40 min at 1,200 rpm. The resin-bound proteins were separated from the unbound fraction by spinning down at 800 g at 4°C. Samples were washed three times with 500 μL of buffer. Results were analysed using tricine or tris-glycine SDS-PAGE and stained with Coomassie brilliant blue. To confirm the successful completion of the phosphorylation reaction, the treated gel was stained using Pro-Q Diamond Phosphoprotein Stain gel (ThermoFisher), according to the manufacturer's protocol, before being stained with Coomassie brilliant blue.

Analytical size exclusion chromatography

Analytical size exclusion chromatography experiments were performed on calibrated Superose 6 Increase 5/150 (Cytiva) or Superdex 200 Increase 5/150 (Cytiva). Purified proteins were mixed in a buffer containing 20 mM HEPES (pH according to the "protein purification" section), 350 mM NaCl, 2.5% glycerol and 1 mM TCEP, with or without the kinase mix. Proteins were diluted to 5 μM, except for the experiments containing MIS18α:MIS18β, which was diluted to 3 μM. For the experiment in **Fig. 3B** and **Fig. S10C**, the proteins were mixed with the following concentrations: MIS18α:MIS18β at 3 µM, MBP-HJURP (394-C or 541-C) at 3 µM and PBD at 13.5 µM (4.5 X molar excess). The mixes were analysed in isocratic conditions at 4°C at a flow rate of 0.15 mL per minute. The fractions were collected and visualised by SDS-PAGE (either tricine or tris-glycine). The successful completion of the phosphorylation reaction was confirmed by staining the gel of the inputs with Pro-Q Diamond Phosphoprotein Stain gel (ThermoFisher), according to the manufacturer's protocol, before being stained with Coomassie brilliant blue.

SEC-MALS

Multi-angle light scattering detector coupled size exclusion chromatography (SEC-MALS) was performed using an Infinity II HPLC (Agilent) coupled to a Superdex 200 10/300 Increase column (GE Healthcare). The column was cooled to 12°C using a Shimadzu column cooler. The MALS system consisted of a Wyatt DAWN 8 MALS detector and a Wyatt Optiplex Refractive Index (RI) detector. The column was pre-equilibrated overnight in a buffer containing 20 mM HEPES pH7.5, 300 mM NaCl and 1 mM TCEP. 50 µL of samples were injected at the following concentrations: 2 mg/mL MBP-M18BP1$^{1-490}$, 2 mg/mL His-PLK1 and a complex of MBP-M18BP1$^{1-490}$ (2 mg/mL) and His-PLK1 (3.8 mg/mL). The theoretical dn/dc for each sample was calculated using SedFit.
The analysis was done using Astra 7.3.2.21, using BSA for peak normalisation. The molar mass for each peak was calculated using the theoretical dn/dc of the proteins and the RI signal. The data was fitted with the Zimm equation.

AlphaFold2 modelling

AlphaFold2 (*42*) was used in version AF2 Multimer 3.2.1 (*49*), which is optimised for predicting complexes between different polymer chains. For the modelling, residues 370-603 of human PLK1 (the PBD domain) and M18BP1 residues 73-96 were used. The protein sequences were taken from UniProt (accession IDs: PLK1 P53550; M18BP1 Q6P0N0) (*50*). For the analysis of the results, see the supplementary text.

Cell culture

Cells were cultured at 37°C in a 5% $CO_2$ atmosphere in DMEM medium (PAN-Biotech) supplemented with 10% tetracycline-free FBS (PAN-Biotech), 50 µg/mL Penicillin/Streptomycin (PAN-Biotech), and 2 mM L-glutamine (PAN-Biotech). For a complete list of the cell lines used for this study see **Table S2**.

Generation of stable cell lines

HeLa Flip-In T-REx stable cell lines were generated by transfecting HeLa CENP-A-SNAP Flp-In T-REx (*17*) with pCDNA5 and pOG44 plasmids according to the protocol previously described (*51*, *52*). For a complete list of the plasmids used for cell line generation see **Table S1**.

CENP-A deposition assay and immunofluorescence

To assess the deposition of the new pool of CENP-A in the early G1 phase, a modified version of a previously published methodology was used (*17*, *23*). Cells were seeded into 12-well dishes and transfected with siRNA oligos using Lipofectamine RNAiMAX Transfection Reagent (Invitrogen), according to the manufacturer's instructions. For a complete list of the plasmid used for cell line generation see **Table S3**. 24 hours after transfection, cells were exposed to media containing 50 ng/mL Doxycycline (Sigma-Aldrich) and 2 mM Thymidine (Sigma-Aldrich), and were incubated for 16 hours to enrich for cells in G1/S state. The following morning, the drugs were washed out and cells were labelled for 30 minutes using 10 mM SNAP-Cell Block (New England Biolabs). After labelling, cells were released in media containing 5 µM S-trityl-L-cysteine (STLC - Sigma-Aldrich) and were incubated for 7h to progress through S, G2 and arrest in mitosis. Then, the newly produced CENP-A-SNAP pool was labelled by exposing the cells to media containing 3 µM SNAP-Cell 647 (New England Biolabs) and 5 µM SLCT for 30 min. Once the labelling was completed, the mitotic cells were collected by shake-off, the drugs were washed out and cells were plated in 24-well dishes containing coverslips. Cells were allowed to attach to the bottom of the wells and exit mitosis for 2.5 hours, then were fixed and immunostained. All the experiments where new CENP-A deposition was not studied followed the same protocol as above but omitted the SNAP-labelling passages.

For the studies using inhibitors (**Figure S1**), the same protocol was used as above with the following modification; after mitotic shake-off and STLC wash-out, cells were released in media containing DMSO or a combination of 300 nM BI2536 (Focus Biomolecules) or 300 nM RO3280 (Tocris Bioscience) and 9 µM RO3306 (Merck Millipore). Cells were released for 2.5 hours before fixation.

Cells were fixed either for 1 minute using ice-cold MeOH and rehydrated with 3 washes for 5 minutes with PBST (PBS supplemented with 0.1% Tween-20), or fixed with PBS + 4% PFA for 10 minutes and permeabilised with PBS + 0.5% Triton-X100 for 15 minutes. Cells were blocked with PBST + 5% BSA for 20 minutes at room temperature and incubated in primary antibodies overnight at 4°C. The following morning, the coverslips were washed 3 times for 5 minutes with PBST, and stained with secondary antibodies for 30 minutes at room temperature. Finally, the coverslips were washed 3 times for 5 minutes with PBST, rinsed quickly in de-ionised water and mounted on microscope slides using Mowiol (EMD Millipore) as the mounting agent. All antibodies were diluted in PBS + 1% BSA. For the complete list of primary and secondary antibodies see **Table S4** and **Table S5**, respectively.

Microscopy

Cells were imaged on a DeltaVision Elite (GE Healthcare) deconvolution microscope, equipped with an IX71 inverted microscope (Olympus, Japan), a UPLSAPO x100/1.40NA oil objective (Olympus) and a pco.edge sCMOS camera (PCO-ECH Inc., USA). Images were acquired as z-sections at 0.2 µm and deconvolved using SoftWoRx (Cytiva).

Data Analysis, Protein Alignment and Figures Generation

Fluorescent signal quantification from fixed cells was performed on maximum intensity projections of single-cell microscope images with FIJI (*53*), using a macro for automated processing. First, kinetochore positions were identified in the FIXME fluorescence channel with a detection scheme based on (*54*), which applies band-pass filtering for contrast enhancement and Otsu thresholding for binary segmentation. Subsequently, mean signal intensities of the respective fluorescence channels were measured from circular regions of interest (ROIs) with a diameter of 7 pixels centred on those positions. Intensities were corrected for local mean background intensity, extracted from 1 pixel-wide ring encircling each ROI.

The raw data from the fluorescent signal quantification was processed as follows. The measurements of each channel were first cleaned from outliers using Prism 8 (GraphPad), using ROUT with a cut-off of Q at 2%. Cleaned data was normalised in Excel software (Microsoft) by dividing each condition by the median value of the control condition (i.e. either the depletion control or the *wild type* sample). Data was visualised as violin plots in Prism 8. For each sample, the median value of each repeat was superimposed to the violin plots as described in the SuperPlots methodology (*55*).

Statistical analysis was performed in Prism 8. All cell biology experiments were repeated at least 3 times to attain statistical significance. We used the Mann-Whitney test for experiments comparing two conditions, and the Kruskall-Wallis test coupled with Dunn's multiple comparisons for experiments comparing 3 or more conditions.

Protein conservation was studied with a Clustal Omega (*56*, *57*) alignment and visualised using Jalview (*58*). All protein sequences used in the alignment were obtained from UniProt (*50*).

The data was placed in Illustrator 2020 (Adobe) to generate the final figures.

### Supplementary Text

AlphaFold2 results

Residues 370-603 of human PLK1 (the PBD domain) and M18BP1 residues 73-96 interacted quite reproducibly (i.e. in 5 models out of 10) and with highly confident pLDDT scores around M18BP1 residues Ser77 and Thr78. The pLDDT scores in the vicinity of M18BP1's Ser93 are only moderately significant in the best of the 10 models, and quite high in the other 9 models. However, the reproducibility of the predicted position of the M18BP1 backbone in this region in 9 out of 10 models indicates that the AlphaFold2 results prediction might still hint at a bona fide interaction in this area next to a positively charged patch of the PBD domain.

**Fig. S1. PLK1 inhibition during mitotic exit prevents new CENP-A deposition.** (**A**) Diagram of the experimental regime in (B-D). (**B**) M18BP1 and new CENP-A levels in human cells where PLK1 was inhibited during mitotic exit using two orthogonal inhibitors (BI2536 and RO3280), alone or in combination with a CDK1 inhibitor (RO3306). (**C**) Quantification of M18BP1 intensity from (B). (**D**) Quantification of new CENP-A intensity from (B). Magenta dots show the median value of each experimental repeat. (**E**) Diagram of the experimental regime in (F-H). (**F**) HJURP-mNG and mScarlet-Mis18α levels in human cells treated during mitotic exit with a PLK1 inhibitor (BI2536) alone or in combination with a CDK1 inhibitor (RO3306). (**G**) Quantification of mScarlet-Mis18α intensity from (F). (**H**) Quantification of HJURP-mNG intensity from (F). Magenta dots show the median value of each experimental repeat.

**Fig. S2. The CENP-A deposition machinery is needed for PLK1 localisation to centromeres in early G1.** (**A**) Diagram of the experimental regime. (**B**) M18BP1, PLK1 and new CENP-A levels in early G1 in human cells treated with siRNA oligos targeting different components of the CENP-A deposition machinery. M18BP1 and the new CENP-A signals were used as a proxy to show the successful depletion of the target proteins. (**C**) Quantification of PLK1 intensity from (B). (**D**) Quantification of M18BP1 intensity from (B). (**E**) Quantification of new CENP-A intensity from (B). Magenta dots show the median value of each experimental repeat.

**Fig. S3. PLK1 binds to the N-terminus of M18BP1 in a phosphorylation-dependent manner.** (**A**) Pull-down experiment showing that PBD binds to a region of M18BP1 encompassing the first 140 residues in a phosphorylation-dependent manner. The success of the kinase reaction was shown using Pro-Q staining. A schematic map of M18BP1 showing the different fragments used in the pull-down experiment of this figure is presented at the top. (**B**) Western blot showing the phosphorylation status of Thr40, Thr78, Ser93 and Ser110 after incubating MBP-M18BP1$^{1\text{-}490}$ with either PLK1 or CDK1. (**C**) Summary diagram of this figure: M18BP1's Thr78 is a quintessential PLK1 site whereas the Ser93 can be phosphorylated by either PLK1 or CDK1.

**Fig. S4. PLK1 is largely sufficient to prime the interaction between M18BP1 and PLK1.** (**A**) Analytical SEC of MBP-M18BP1$^{1\text{-}490}$ incubated with MBP-PBD in the presence or absence of PLK1 activity. AURORA A and BORA control show that the interaction is specific to PLK1 activity, and not promoted by the activation mix of PLK1. (**B**) Analytical SEC of MBP-M18BP1$^{1\text{-}490}$ incubated with MBP-PBD in the presence or absence of CDK1 activity. (**C**) Coomassie and Pro-Q staining of the inputs of the analytical SEC from (A).

**Fig. S5. Depletion controls for the experiment in Fig. 1F-H.** (**A**) Diagram of the experimental regime. (**B**) Endogenous M18BP1 levels in the parental cells show the depletion extent of the protein across each repeat of this experiment. (**C**) Quantification of the endogenous M18BP1 intensity from (B). Magenta dots show the median value of each experimental repeat.

**Fig. S6. M18BP1 and PLK1 bind with a 1:1 stoichiometry.** (**A**) Coomassie and Pro-Q staining of the inputs of the SEC-MALS from (B). (**B**) Chromatogram of the SEC-MALS measurement showing the calculated molecular weight of each sample injected. (**C**) Plot of the predicted alignment error of the AlfaFold2 modelling from Fig. 1I.

**Fig. S7. M18BP1 mutants and MIS18α can localise to centromeres in early G1 (quantification from Fig. 2A-B).** (**A**) Diagram of the experimental regime. (**B**) Quantification of the endogenous M18BP1 intensity from the parental cells from Fig. 2A. (**C**) Quantification of EGFP-M18BP1 intensity from Fig. 2A. (**D**) Quantification of mCherry-MIS18α intensity from Fig. 2A. Magenta dots show the median value of each experimental repeat.

**Fig. S8. M18BP1 point mutants bind to MIS18αβ similarly to the *wild-type.*** Analytical SEC of MBP-M18BP1$^{1\text{-}490}$ (*wild-type* or mutants) incubated with MIS18α:mScarlet-I-MIS18β complex. A cartoon of the question addressed by the experiment is displayed at the top.

**Fig. S9. Depletion controls from Fig. 2C-D.** (**A**) Diagram of the experimental regime. (**B**) Quantification of the endogenous M18BP1 levels from the parental cells shows the depletion extent of the protein across each repeat of this experiment. (**C**) Quantification of the endogenous M18BP1 intensity from (B). Magenta dots show the median value of each experimental repeat.

**Fig. S10. PLK1 activity and its binding to MIS18α enhances HJURP affinity for MIS18αβ.** (**A**) Coomassie and Pro-Q staining of the inputs of the analytical SEC from Fig. 3B. (**B**) Coomassie and Pro-Q staining of the inputs of the analytical SEC from (C). (**C**) Analytical SEC shows that MIS18αβ and MBP-HJURP$^{394\text{-}540}$ can bind in the presence of PBD and PLK1 phosphorylation.

**Fig. S11. PLK1 binds to MIS18α N-terminus in a phosphorylation-dependent manner.** (**A**) Analytical SEC shows that MIS18αβ and MBP-PBD bind in the presence of PLK1 phosphorylation. AURORA A and BORA control show that the interaction is specific to PLK1 activity, and not promoted by the activation mix of PLK1. (**B**) Analytical SEC shows that MIS18α$^{S53A\text{-}S54A}$β and MBP-PBD do not bind even in the presence of PLK1 phosphorylation. Cartoons of the experimental question are above each experiment.

**Fig. S12. PLK1 binds to HJURP's C-terminus in a phosphorylation-dependent manner.** (**A**) Analytical SEC of MBP-HJURP$^{394\text{-}C}$ incubated with MBP-PBD in the presence or absence of PLK1 activity. AURORA A and BORA control show that the interaction is specific to PLK1 activity, and not promoted by the activation mix of PLK1. (**B**) Pull-down experiment showing that PBD binds to a region of HJURP encompassing the last 207 residues in a phosphorylation-dependent manner. The success of the kinase reaction was shown using Pro-Q staining. A schematic map of HJRP showing the different fragments used in the pull-down experiment of this figure is shown at the top. (**C**) Pull-down experiment shows that PBD binds to HJURPS's Ser653-Thr654 motif. The success of the kinase reaction was shown using Pro-Q staining. (**D**) Analytical SEC of MBP-HJURP$^{394\text{-}C}$ (*wild-type* or mutant) incubated with MBP-PBD in the presence or absence of PLK1 activity. Cartoons of the experimental question are above or next to each experiment.

**Fig. S13. PLK1 binding to both MIS18α and HJURP is essential for the MIS18α:HJURP complex formation.** (**A**) Cartoon of the question addressed by this experiment. (**B**) Coomassie and Pro-Q staining of the inputs of the analytical SEC from (C-F). (**C-F**) Analytical SEC shows that point mutants of MIS18αβ and HJURP$^{541\text{-}C}$ cannot bind to each other any longer in the presence of PBD and PLK1 phosphorylation. This is the complete experiment from Fig. 3 E.

**Fig. S14. Depletion controls for Fig. 3F-H.** (**A**) Diagram of the experimental regime. (**B**) Endogenous M18BP1 and new CENP-A levels in the parental cells show the depletion extent of the protein across each repeat of the experiment. (**C**) Quantification of the endogenous M18BP1 intensity from (B). (**D**) Quantification of new CENP-A intensity from (B). (**E**) Quantification of mCherry-MIS18α intensity from Fig. 3F. Magenta dots show the median value of each experimental repeat.

**Fig. S15. Preventing PLK1 binding to MIS18α or HJURP mildly affects new CENP-A deposition.** (**A**) Diagram of the experimental regime. (**B**) EGFP-M18BP1, mCherry-MIS18α and new CENP-A levels at centromeres in human cells expressing MIS18α (*wild-type* or mutant) in early G1 phase. The upper panels show the parental cells as controls for the endogenous M18BP1 depletion. (**C**) Quantification of the new CENP-A intensity from (B). (**D**) Quantification of the endogenous M18BP1 intensity from (B). (**E**) Quantification of EGFP-M18BP1 intensity from (B). (**F**) Quantification of mCherry-MIS18α intensity from (B). Magenta dots show the median value of each experimental repeat. (**G**) Endogenous M18BP1 and new CENP-A levels in the parental cells show the depletion extent of the protein across each repeat of the experiment in panels I-L. (**H**) Quantification of new CENP-A intensity from (G). (**I**) mCherry-M18BP1, mNeonGreen-HJURP and new CENP-A levels at centromeres in human cells expressing HJURP (*wild-type* or mutant) in early G1 phase. (**J**) Quantification of new CENP-A intensity from (I). (**K**) Quantification of HJURP-mNG intensity from (I). (**L**) Quantification of mCherry-M18BP1 intensity from (I). Magenta dots show the median value of each experimental repeat.

**Fig. S16. Depletion controls for Fig. 4A-B.** (**A**) Diagram of the experimental regime. (**B**) Endogenous M18BP1 levels in the parental cells show the depletion extent of the protein across each repeat of the experiment. (**C**) Quantification of the endogenous M18BP1 intensity from (B). (**D**) Quantification of new CENP-A intensity from (B). (**E**) Quantification of EGFP-M18BP1 intensity from Fig. 4A. (**F**) Quantification of mCherry-MIS18α intensity from Fig. 4A. Magenta dots show the median value of each experimental repeat.

**Fig. S17. M18BP1$^{56\text{-}98}$ is a minimal PLK1-binding fragment.** (**A**) Cartoon showing the location of the minimal PLK1-binding site on M18BP1 designed for this experiment. The inset below displays the evolutionary conservation of the domain. (**B**) Analytical SEC of MBP-M18BP1$^{56\text{-}98}$ incubated with a stoichiometric amount of His-PLK1 in the presence or absence of kinase activity.

**Fig. S18. Grafting M18BP1's minimal PLK1-binding fragment onto HJURP does not sustain new CENP-A deposition in the absence of PLK1 at centromeres.** (**A**) Cartoon showing the grafting strategy to generate a chimeric HJURP bearing M18BP1's PLK1-binding site. (**B**) Diagram of the experimental regime. (**C**) mCherry-M18BP1, HJURP-mNG and new CENP-A levels at centromeres in human cells co-expressing in different combinations M18BP1 (*wild-type* or S77A-T78V-S93A mutant) and HJURP (*wild-type* or the graft mutant) in early G1 phase. The upper panels display the parental cells as controls for the endogenous M18BP1 depletion. (**D**) Quantification of new CENP-A intensity from (C). (**E**) Quantification of mCherry-M18BP1 intensity from (C). (**F**) Quantification of HJURP-mNG intensity from (C). (**G**) Quantification of the endogenous M18BP1 intensity from (C). Magenta dots show the median value of each experimental repeat.

**Fig. S19. Centromeric localisation of PLK1 and MIS18α in early G1 is not altered in cells expressing MIS18α$^{S53A\text{-}S54A}$ mutants.** (**A**) Diagram of the experimental regime. (**B**) Endogenous M18BP1 levels in the parental cells show the depletion extent of the protein across each repeat of the experiment in panels D-G. (**C**) Quantification of the endogenous M18BP1 intensity from (B). (**D**) mScarlet-MIS18α, mNG-PLK1 and M18BP1 pThr702 levels in human cells expressing mScarlet-MIS18α (*wild-type* or the S53A-S54A mutant) in early G1 phase. (**E**) Quantification of mScarlet-MIS18α intensity from (D). (**F**) Quantification of mNG-PLK1 intensity from (D). (**G**) Quantification of M18BP1 pThr702 intensity from (D). Magenta dots show the median value of each experimental repeat.

**Fig. S20. The PLK1-docking sites on M18BP1, MIS18α and HJURP are close to their interaction domains.** Cartoon summarising the PLK1-binding sites discovered in this study. PLK1 binds to M18BP1, MIS18α and HJURP in consensus sites close to the domains used to assemble the complete CENP-A deposition machinery.

**Table S1.**

List of the plasmids used in this study.

| Expressed Construct (backbone) | Purpose | Source |
|---|---|---|
| (pGEX6PT) GST | protein expression | *Pan et al.*, 2017 |
| (pETDuet) MBP-8His | protein expression | *Pan et al.*, 2017 |
| (pGEX6PT) GST-M18BP1$^{1\text{-}490}$ | protein expression | This study |

| **Expressed Construct (backbone)** | **Purpose** | **Source** |
|---|---|---|
| (pGEX6PT) GST-M18BP1$^{1\text{-}140}$ | protein expression | This study |
| (pGEX6PT) GST-M18BP1$^{141\text{-}311}$ | protein expression | This study |
| (pGEX6PT) GST-M18BP1$^{312\text{-}490}$ | protein expression | This study |
| (pETDuet) MBP-M18BP1$^{1\text{-}490}$-8His | protein expression | *Pan et al.*, 2017 |
| (pETDuet) MBP-M18BP1$^{1\text{-}490;\ T78V}$-8His | protein expression | This study |
| (pETDuet) MBP-M18BP1$^{1\text{-}490;\ S77A\text{-}T78V}$-8His | protein expression | This study |
| (pETDuet) MBP-M18BP1$^{1\text{-}490;\ S93A}$-8His | protein expression | This study |
| (pETDuet) MBP-M18BP1$^{1\text{-}490;\ S77A\text{-}T78V\text{-}S93A}$-8His | protein expression | This study |
| (pETDuet) 6His-MBP-M18BP1$^{56\text{-}98}$ | protein expression | This study |
| (pETDuet) 6His-MIS18α; MBP-MIS18β | protein expression | *Pan et al.*, 2017 |
| (pETDuet) 6His-mScarlet-I-MIS18β; MIS18α | protein expression | This study |
| (pETDuet) 6His-MIS18α$^{56\text{-}C}$; MBP-MIS18β | protein expression | *Pan et al.*, 2019 |
| (pETDuet) 6His-MIS18α$^{S53A\text{-}S54A}$; MBP-MIS18β | protein expression | This study |
| (pETDuet) MBP-HJURP$^{1\text{-}393}$-8His | protein expression | *Pan et al.*, 2019 |
| (pETDuet) MBP-HJURP$^{394\text{-}748}$-8His | protein expression | *Pan et al.*, 2019 |
| (pETDuet) MBP-HJURP$^{394\text{-}540}$-8His | protein expression | *Pan et al.*, 2019 |
| (pETDuet) MBP-HJURP$^{541\text{-}748}$-8His | protein expression | *Pan et al.*, 2019 |

| Expressed Construct (backbone) | Purpose | Source |
|---|---|---|
| (pETDuet) MBP-HJURP$^{541\text{-}748;\ S548A\text{-}S549A}$-8His | protein expression | This study |
| (pETDuet) MBP-HJURP$^{541\text{-}748;\ S641A\text{-}S642A}$-8His | protein expression | This study |
| (pETDuet) MBP-HJURP$^{541\text{-}748;\ S653A\text{-}T654V}$-8His | protein expression | This study |
| (pETDuet) MBP-HJURP$^{394\text{-}748;\ S653A\text{-}T654V}$-8His | protein expression | This study |
| (pLIB) 6His-PLK1 | protein expression | *Singh, Pesenti et al.*, 2020 |
| (pETDuet) 6His-MBP-PBD | protein expression | *Singh, Pesenti et al.*, 2020 |
| (pLIB) GST-CDK1 | protein expression | *Huis in't Veld et al.*, 2021 |
| (pLIB) His-TEV-CKS1 | protein expression | *Huis in't Veld et al.*, 2021 |
| (pLIB) His-TEV-CyclinB1 | protein expression | *Huis in't Veld et al.*, 2021 |
| (pLIB) MBP-CIV1 | protein expression | *Huis in't Veld et al.*, 2021 |
| (pET28) 6His-AURORA A | protein expression | This study |
| (pETDuet) MBP-BORA$^{1\text{-}224}$-8His | protein expression | This study |
| (pCDNA5) mNeonGreen-PLK1-P2AT2A-mScarlet-M18BP1 | cell line generation | This study |
| (pCDNA5) mNeonGreen-PLK1-P2AT2A-mScarlet-M18BP1$^{S77A\text{-}T78V}$ | cell line generation | This study |
| (pCDNA5) mNeonGreen-PLK1-P2AT2A-mScarlet-M18BP1$^{S93A}$ | cell line generation | This study |
| (pCDNA5) mNeonGreen-PLK1-P2AT2A-mScarlet-M18BP1$^{S77A\text{-}T78V\text{-}S93A}$ | cell line generation | This study |
| (pCDNA5) mNeonGreen-PLK1-P2AT2A-mScarlet-MIS18α | cell line generation | This study |

| Expressed Construct (backbone) | Purpose | Source |
|---|---|---|
| (pCDNA5) mNeonGreen-PLK1-P2AT2A-mScarlet-MIS18α$^{S53A-S54A}$ | cell line generation | This study |
| (pCDNA5) EGFP-M18BP1-P2AT2A-mCherry-MIS18α | cell line generation | *Pan et al.*, 2019 |
| (pCDNA5) EGFP-M18BP1$^{S77A-T78V}$-P2AT2A-mCherry-MIS18α | cell line generation | This study |
| (pCDNA5) EGFP-M18BP1$^{S93A}$-P2AT2A-mCherry-MIS18α | cell line generation | This study |
| (pCDNA5) EGFP-M18BP1$^{S77A-T78V-S93A}$-P2AT2A-mCherry-MIS18α | cell line generation | This study |
| (pCDNA5) EGFP-M18BP1-P2AT2A-mCherry-MIS18α$^{S53A-S54A}$ | cell line generation | This study |
| (pCDNA5) M18BP1-EGFP-P2AT2A-mCherry-MIS18α | cell line generation | This study |
| (pCDNA5) M18BP1$^{S77A-T78V-S93A}$-EGFP-P2AT2A-mCherry-MIS18α | cell line generation | This study |
| (pCDNA5) M18BP1-EGFP-P2AT2A-mCherry-MIS18α$^{56-C}$ | cell line generation | This study |
| (pCDNA5) M18BP1$^{S77A-T78V-S93A}$-EGFP-P2AT2A-mCherry-MIS18α$^{56-C}$ | cell line generation | This study |
| (pCDNA5) HJURP-mNeonGreen-P2AT2A-mScarlet-M18BP1 | cell line generation | This study |
| (pCDNA5) HJURP-mNeonGreen-P2AT2A-mScarlet-M18BP1$^{S77A-T78V-S93A}$ | cell line generation | This study |
| (pCDNA5) HJURP-mNeonGreen-P2AT2A-mCherry-M18BP1 | cell line generation | This study |
| (pCDNA5) HJURP-mNeonGreen-P2AT2A-mCherry-M18BP1$^{S77A-T78V-S93A}$ | cell line generation | This study |
| (pCDNA5) HJURP$^{S653A-T654V}$-mNeonGreen-P2AT2A-mCherry-M18BP1 | cell line generation | This study |
| (pCDNA5) HJURP$^{graft}$-mNeonGreen-P2AT2A-mCherry-M18BP1$^{S77A-T78V-S93A}$ | cell line generation | This study |
| (pCDNA5) HJURP-mNeonGreen-P2AT2A-mCherry-MIS18α | cell line generation | This study |

| Expressed Construct (backbone) | Purpose | Source |
|---|---|---|
| (pCDNA5) HJURP$^{S653A\text{-}T654V}$-mNeonGreen-P2AT2A-mCherry-MIS18α$^{S53A\text{-}S54A}$ | cell line generation | This study |
| (pCDNA5) HJURP-mNeonGreen-P2AT2A-mScarlet-MIS18α | cell line generation | This study |
| pOG44 | cell line generation | Donated by Stephen Taylor |

**Table S2.**

List of the cell lines used in this study.

| Cell line name | Source |
|---|---|
| HeLa Flp-In T-REx CENP-A-SNAP | *Pan et al.*, 2017 |
| HeLa CENP-A-SNAP; mNeonGreen-PLK1-P2AT2A-mScarlet-M18BP1 | This study |
| HeLa CENP-A-SNAP; mNeonGreen-PLK1-P2AT2A-mScarlet-M18BP1$^{S77A\text{-}T78V}$ | This study |
| HeLa CENP-A-SNAP; mNeonGreen-PLK1-P2AT2A-mScarlet-M18BP1$^{S93A}$ | This study |
| HeLa CENP-A-SNAP; mNeonGreen-PLK1-P2AT2A-mScarlet-M18BP1$^{S77A\text{-}T78V\text{-}S93A}$ | This study |
| HeLa CENP-A-SNAP; mNeonGreen-PLK1-P2AT2A-mScarlet-MIS18α | This study |
| HeLa CENP-A-SNAP; mNeonGreen-PLK1-P2AT2A-mScarlet-MIS18α$^{S53A\text{-}S54A}$ | This study |
| HeLa CENP-A-SNAP; EGFP-M18BP1-P2AT2A-mCherry-MIS18α | *Pan et al.*, 2017 |
| HeLa CENP-A-SNAP; EGFP-M18BP1$^{S77A\text{-}T78V}$-P2AT2A-mCherry-MIS18α | This study |
| HeLa CENP-A-SNAP; EGFP-M18BP1$^{S93A}$-P2AT2A-mCherry-MIS18α | This study |
| HeLa CENP-A-SNAP; EGFP-M18BP1$^{S77A\text{-}T78V\text{-}S93A}$-P2AT2A-mCherry-MIS18α | This study |
| HeLa CENP-A-SNAP; EGFP-M18BP1-P2AT2A-mCherry-MIS18α$^{S53A\text{-}S54A}$ | This study |
| HeLa CENP-A-SNAP; M18BP1-EGFP-P2AT2A-mCherry-MIS18α | This study |
| HeLa CENP-A-SNAP; M18BP1$^{S77A\text{-}T78V\text{-}S93A}$-EGFP-P2AT2A-mCherry-MIS18α | This study |
| HeLa CENP-A-SNAP; M18BP1-EGFP-P2AT2A-mCherry-MIS18α$^{56\text{-}C}$ | This study |
| HeLa CENP-A-SNAP; M18BP1$^{S77A\text{-}T78V\text{-}S93A}$-EGFP-P2AT2A-mCherry-MIS18α$^{56\text{-}C}$ | This study |
| HeLa CENP-A-SNAP; HJURP-mNeonGreen-P2AT2A-mScarlet-M18BP1 | This study |
| HeLa CENP-A-SNAP; HJURP-mNeonGreen-P2AT2A-mScarlet-M18BP1$^{S77A\text{-}T78V\text{-}S93A}$ | This study |

| Cell line name | Source |
|---|---|
| HeLa CENP-A-SNAP; HJURP-mNeonGreen-P2AT2A-mCherry-M18BP1 | This study |
| HeLa CENP-A-SNAP; HJURP-mNeonGreen-P2AT2A-mCherry-M18BP1$^{S77A-T78V-S93A}$ | This study |
| HeLa CENP-A-SNAP; HJURP$^{S653A-T654V}$-mNeonGreen-P2AT2A-mCherry-M18BP1 | This study |
| HeLa CENP-A-SNAP; HJURP$^{graft}$-mNeonGreen-P2AT2A-mCherry-M18BP1$^{S77A-T78V-S93A}$ | This study |
| HeLa CENP-A-SNAP; HJURP-mNeonGreen-P2AT2A-mCherry-MIS18α | This study |
| HeLa CENP-A-SNAP; HJURP$^{S653A-T654V}$-mNeonGreen-P2AT2A-mCherry-MIS18α$^{S53A-S54A}$ | This study |
| HeLa CENP-A-SNAP; HJURP-mNeonGreen-P2AT2A-mScarlet-MIS18α | This study |

**Table S3.**

List of the siRNA oligos used in this study. All the oligos were provided by Sigma-Aldrich.

| Target mRNA | Sequence (3'-5') | Concentration | Source |
|---|---|---|---|
| M18BP1 | GAAGUCUGGUGUUAGGAAAdTdT | 10 nM | *Fujita et al.*, 2007 |
| MIS18α | CAGAAGCUAUCCAAACGUGdTdT | 10 nM | *Fujita et al.*, 2007 |
| HJURP | GGAGUGUUAUUUAUCUUCCCACUUG | 10 nM | *Pan et al.*, 2019 |

**Table S4.**

List of the primary antibodies used in this study.

| Epitope | Species | Manufacturer (cat. number) |
|---|---|---|
| α-Tubulin | Mouse monoclonal | Sigma-Aldrich (T9026) |
| CREST | Human autoimmune serum | Antibodies Inc. (15-234) |
| M18BP1 | Rat polyclonal | This study (n.38) |
| M18BP1 pT40 | Mouse monoclonal | This study (n. GT72) |
| M18BP1 pT78 | Mouse monoclonal | This study (n. GU64) |
| M18BP1 pS93 | Mouse monoclonal | This study (n. GW03) |

| Epitope | Species | Manufacturer (cat. number) |
|---|---|---|
| M18BP1 pS110 | Mouse monoclonal | This study (n. GX09) |
| M18BP1 pT702 | Rabbit polyclonal | Merck Millipore (ABE1847) |
| MBP | Mouse monoclonal | New England Biosciences (E8032S) |
| PLK1 | Mouse monoclonal | Abcam (ab17057) |

**Table S5.**

List of the secondary antibodies used in this study.

| Name | Species | Manufacturer (cat. number) |
|---|---|---|
| α-Human DyLight 405 | Donkey | Jackson Immuno Research (709-475-149) |
| α-Mouse Rodamine Red | Goat | Jackson Immuno Research (115-295-0030) |
| α-Rabbit Alexa Fluor 647 | Donkey | Jackson Immuno Research (711-606-152) |
| α-Rat Alexa Fluor 488 | Donkey | Jackson Immuno Research (712-545-153) |

**References:**

48. P. J. Huis In ’t Veld, S. Wohlgemuth, C. Koerner, F. Müller, P. Janning, A. Musacchio, Reconstitution and use of highly active human CDK1:Cyclin-B:CKS1 complexes. *Protein Sci.* **31**, 528–537 (2022).

49. R. Evans, M. O’Neill, A. Pritzel, N. Antropova, A. Senior, T. Green, A. Žídek, R. Bates, S. Blackwell, J. Yim, O. Ronneberger, S. Bodenstein, M. Zielinski, A. Bridgland, A. Potapenko, A. Cowie, K. Tunyasuvunakool, R. Jain, E. Clancy, P. Kohli, J. Jumper, D. Hassabis, Protein complex prediction with AlphaFold-Multimer. *bioRxiv* (2022), p. 2021.10.04.463034.

50. UniProt Consortium, UniProt: the Universal Protein Knowledgebase in 2023. *Nucleic Acids Res.* **51**, D523–D531 (2023).

51. A. Tighe, V. L. Johnson, S. S. Taylor, Truncating APC mutations have dominant effects on proliferation, spindle checkpoint control, survival and chromosome stability. *J. Cell Sci.* **117**, 6339–6353 (2004).

52. A. Tighe, O. Staples, S. Taylor, Mps1 kinase activity restrains anaphase during an unperturbed mitosis and targets Mad2 to kinetochores. *J. Cell Biol.* **181**, 893–901 (2008).

53. J. Schindelin, I. Arganda-Carreras, E. Frise, V. Kaynig, M. Longair, T. Pietzsch, S. Preibisch, C. Rueden, S. Saalfeld, B. Schmid, J.-Y. Tinevez, D. J. White, V. Hartenstein, K. Eliceiri, P. Tomancak, A. Cardona, Fiji: an open-source platform for biological-image analysis. *Nat. Methods*. **9**, 676 (2012).

54. D. L. Bodor, M. G. Rodríguez, N. Moreno, L. E. T. Jansen, Analysis of protein turnover by quantitative SNAP-based pulse-chase imaging. *Curr. Protoc. Cell Biol.* **Chapter 8**, Unit8.8 (2012).

55. S. J. Lord, K. B. Velle, R. D. Mullins, L. K. Fritz-Laylin, SuperPlots: Communicating reproducibility and variability in cell biology. *J. Cell Biol.* **219** (2020), doi:10.1083/jcb.202001064.

56. M. Goujon, H. McWilliam, W. Li, F. Valentin, S. Squizzato, J. Paern, R. Lopez, A new bioinformatics analysis tools framework at EMBL-EBI. *Nucleic Acids Res.* **38**, W695–W699 (2010).

57. F. Sievers, A. Wilm, D. Dineen, T. J. Gibson, K. Karplus, W. Li, R. Lopez, H. McWilliam, M. Remmert, J. S. O. Ding, J. D. Thompson, D. G. Higgins, Fast, scalable generation of high-quality protein multiple sequence alignments using Clustal Omega. *Mol. Syst. Biol.* **7**, 1–6 (2011).

58. A. M. Waterhouse, J. B. Procter, D. M. A. Martin, M. Clamp, G. J. Barton, Jalview Version 2--a multiple sequence alignment editor and analysis workbench. *Bioinformatics*. **25**, 1189–1191 (2009).

**Figure S1.** Plk1 inhibition during mitotic exit prevents new CENP-A deposition.

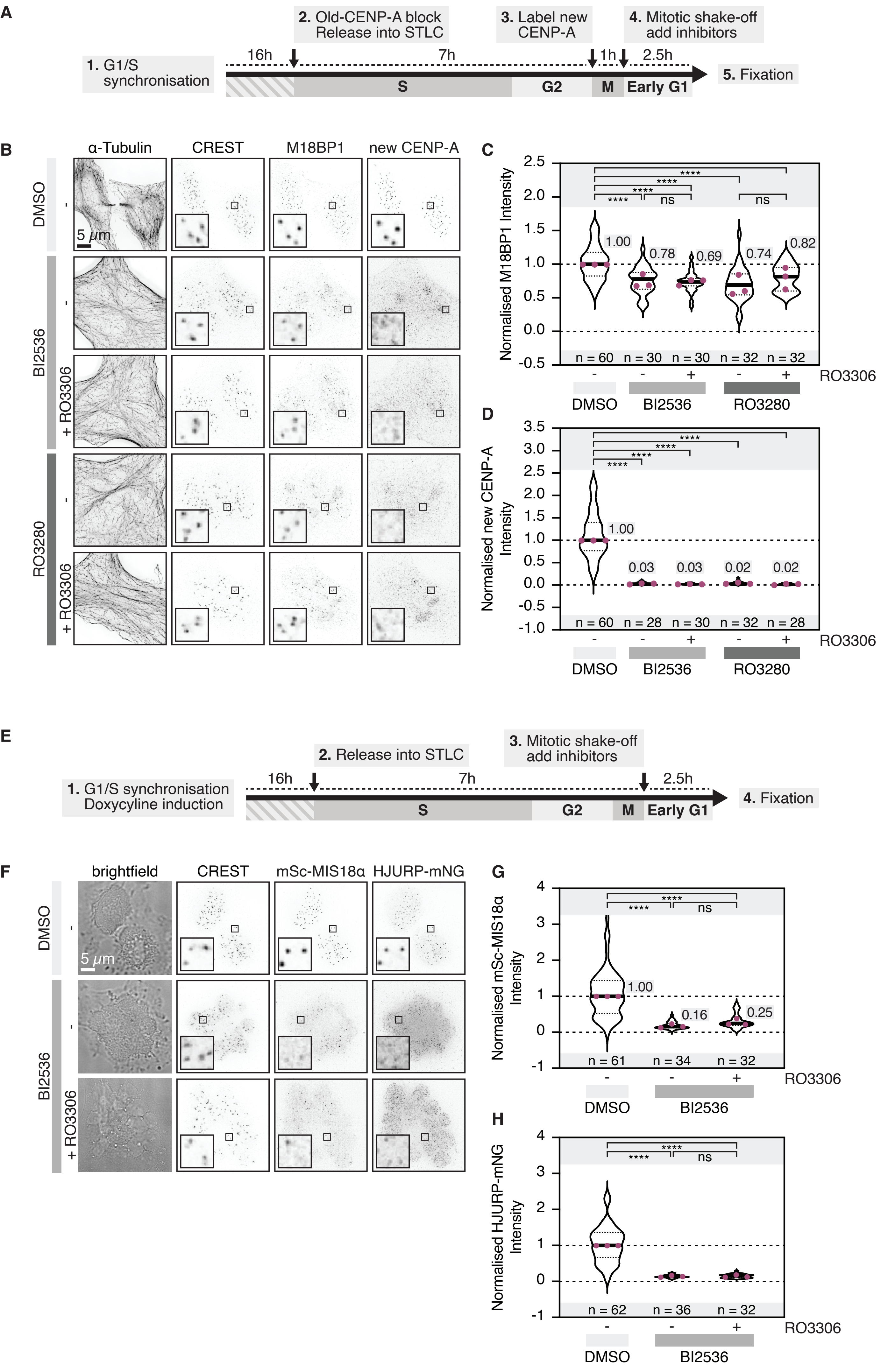

**Figure S2.** The CENP-A deposition machinery is needed for PLK1 localisation to centromeres in early G1.

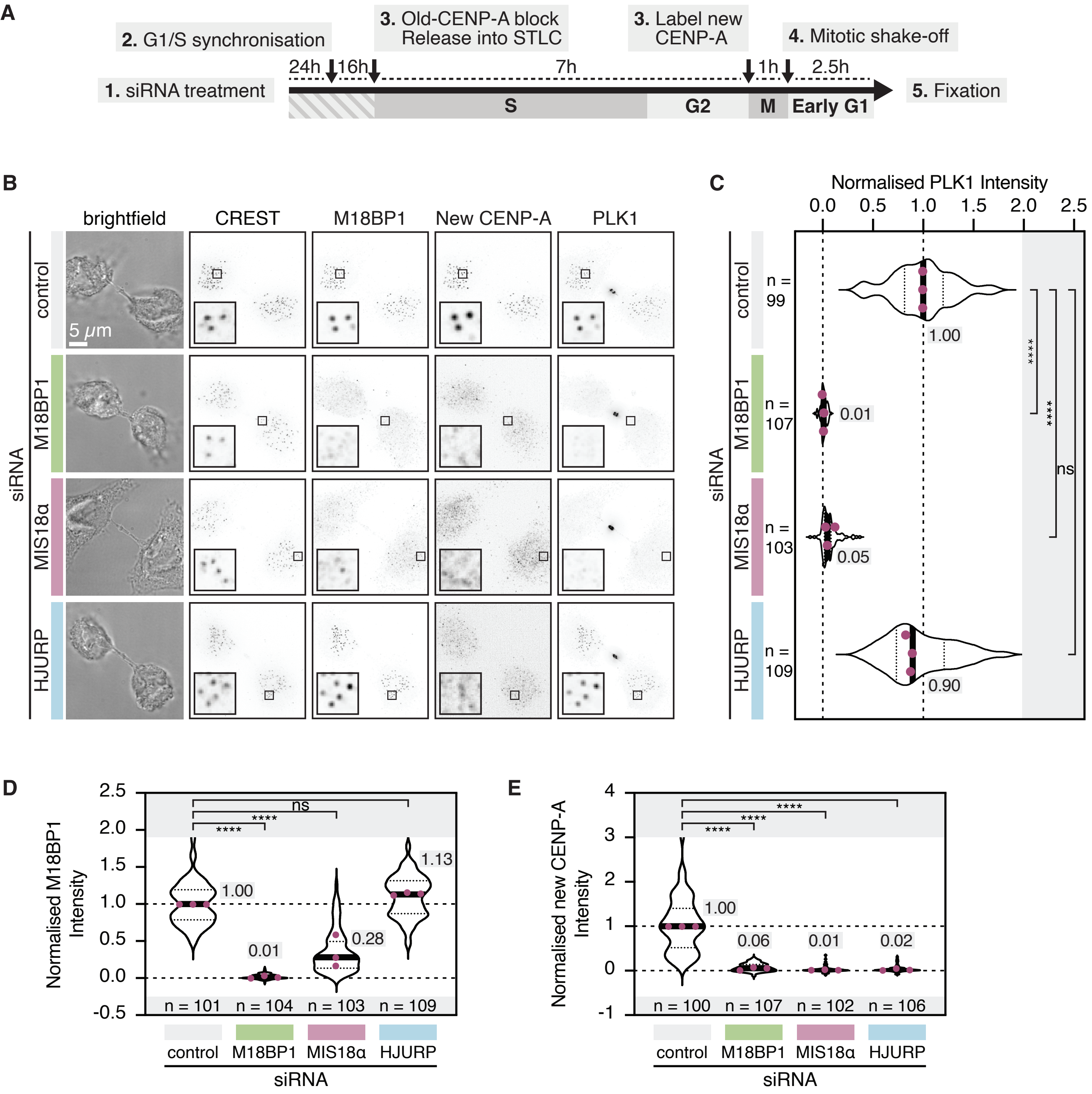

**Figure S3.** PLK1 binds to the N-terminus of M18BP1 in a phosphorylation-dependent manner.

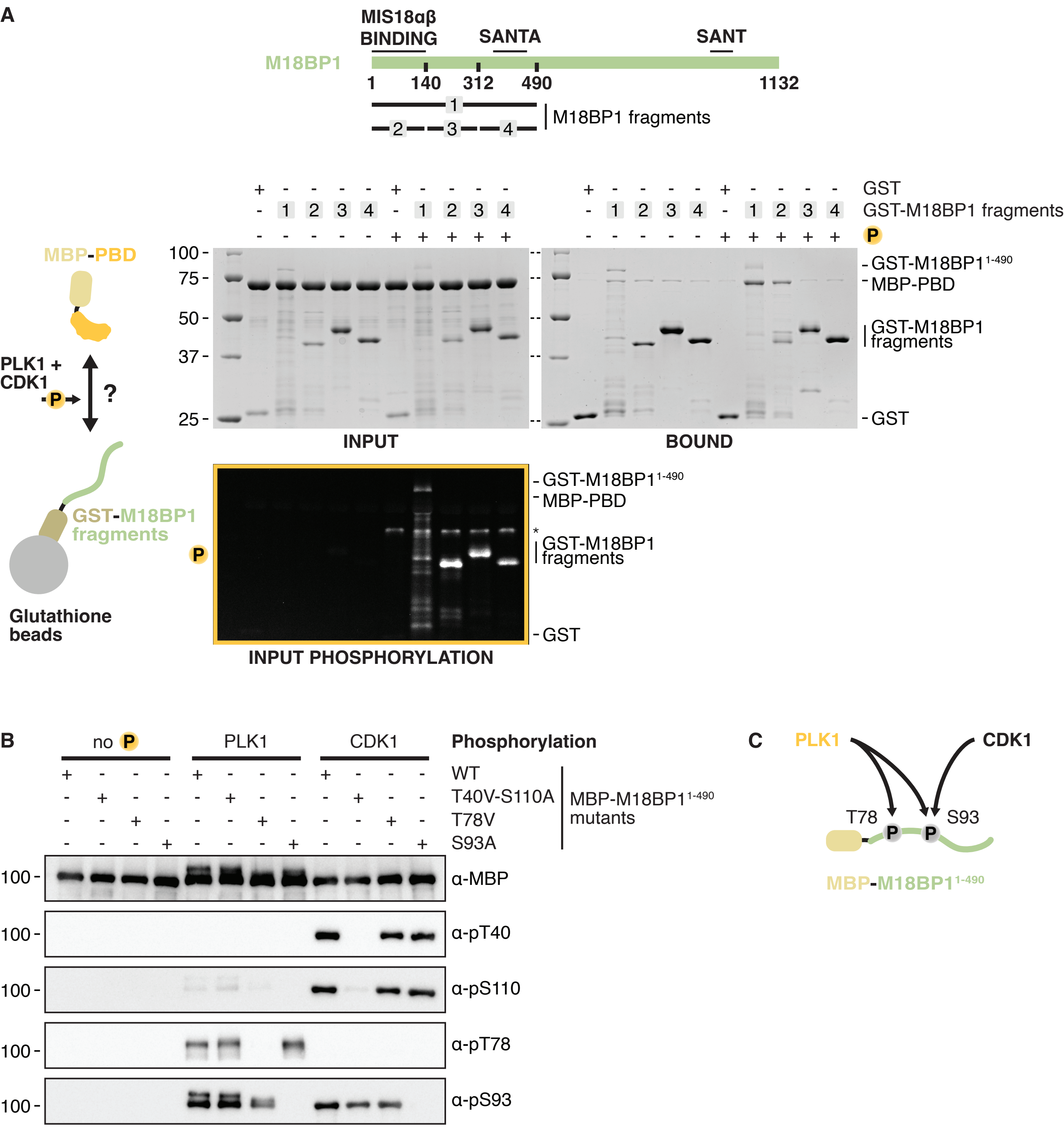

**Figure S4.** PLK1 is largely sufficient to prime the interaction between M18BP1 and PLK1.

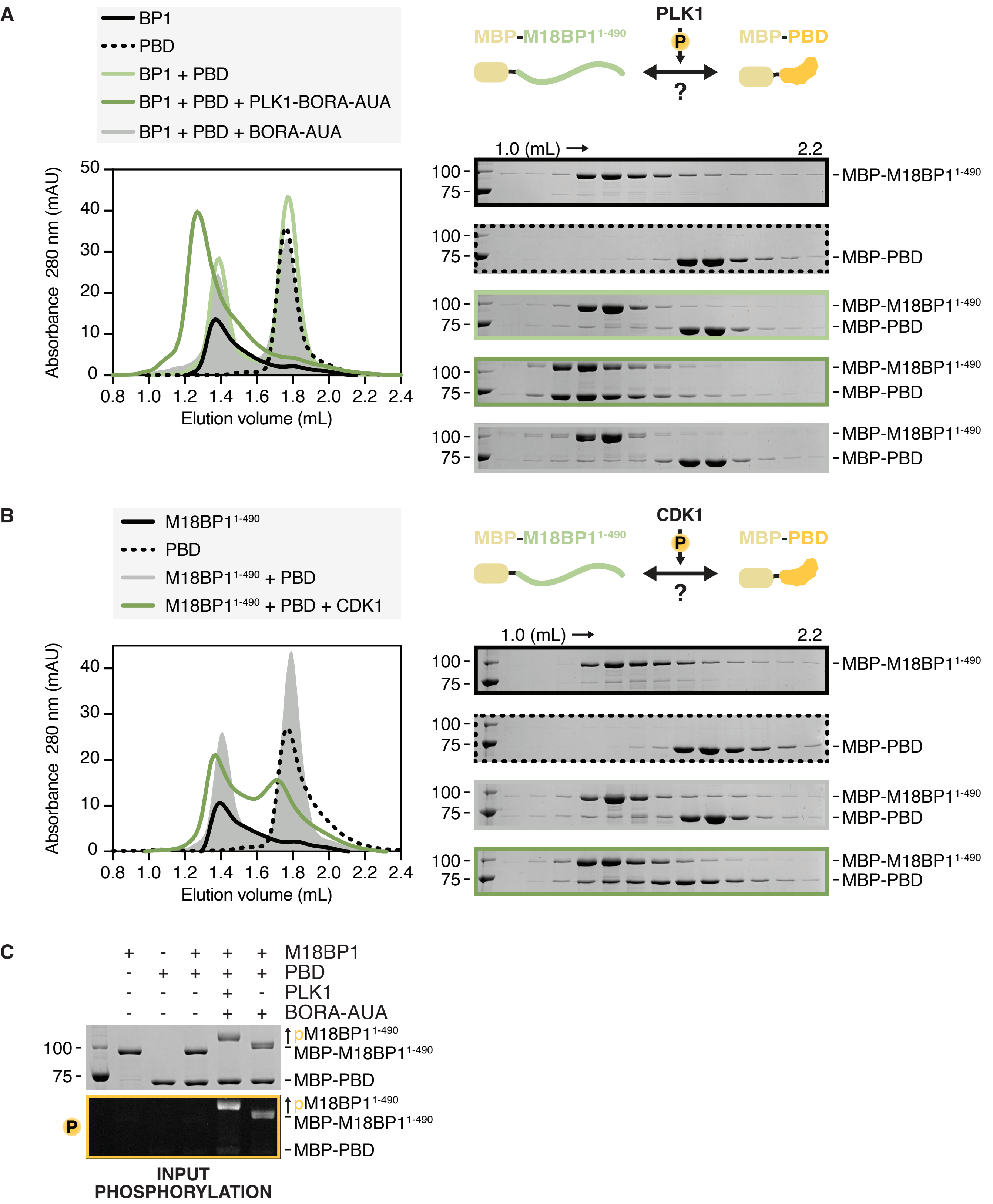

**Figure S5.** Depletion controls for the experiment in Fig. 1F-H.

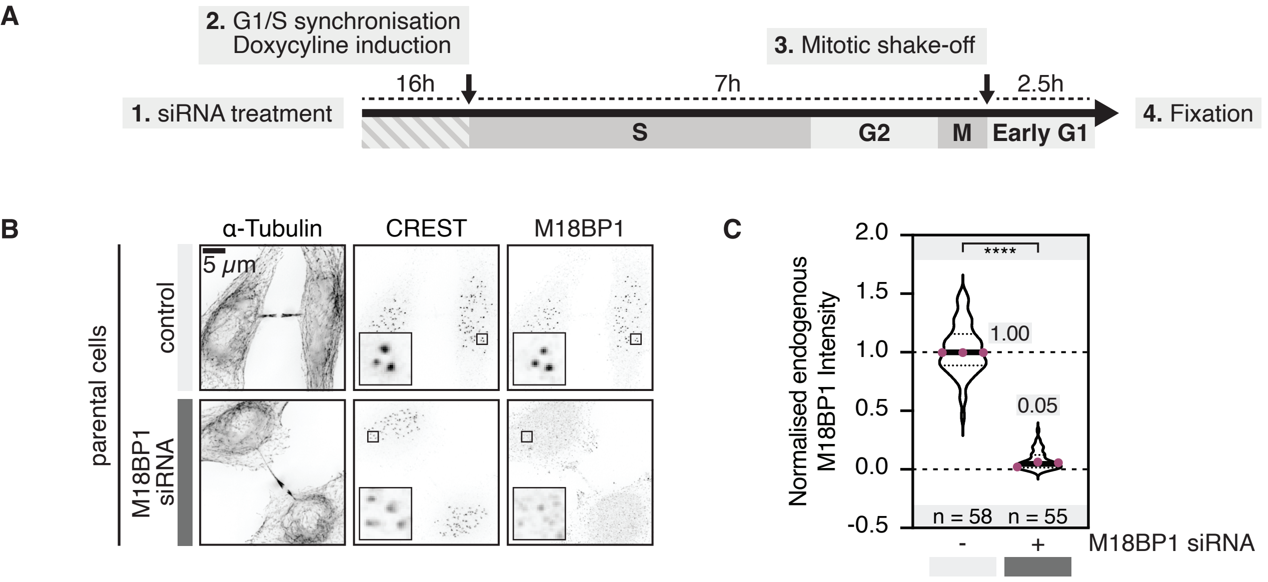

**Figure S6.** M18BP1 and PLK1 bind with a 1:1 stoichiometry.

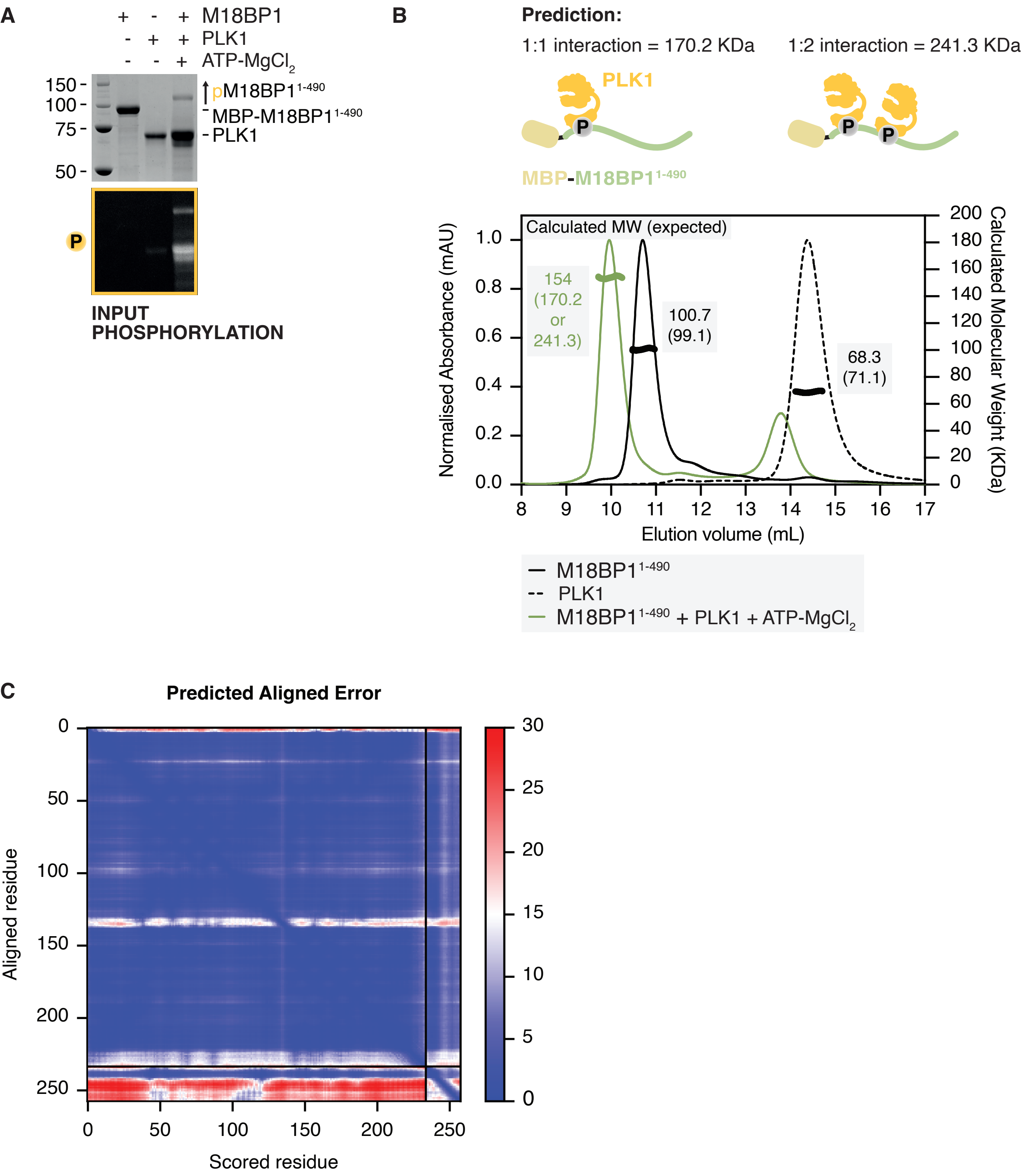

**Figure S7.** M18BP1 mutants and MIS18α can localise to centromeres in early G1 (quantification from Fig. 2A-B).

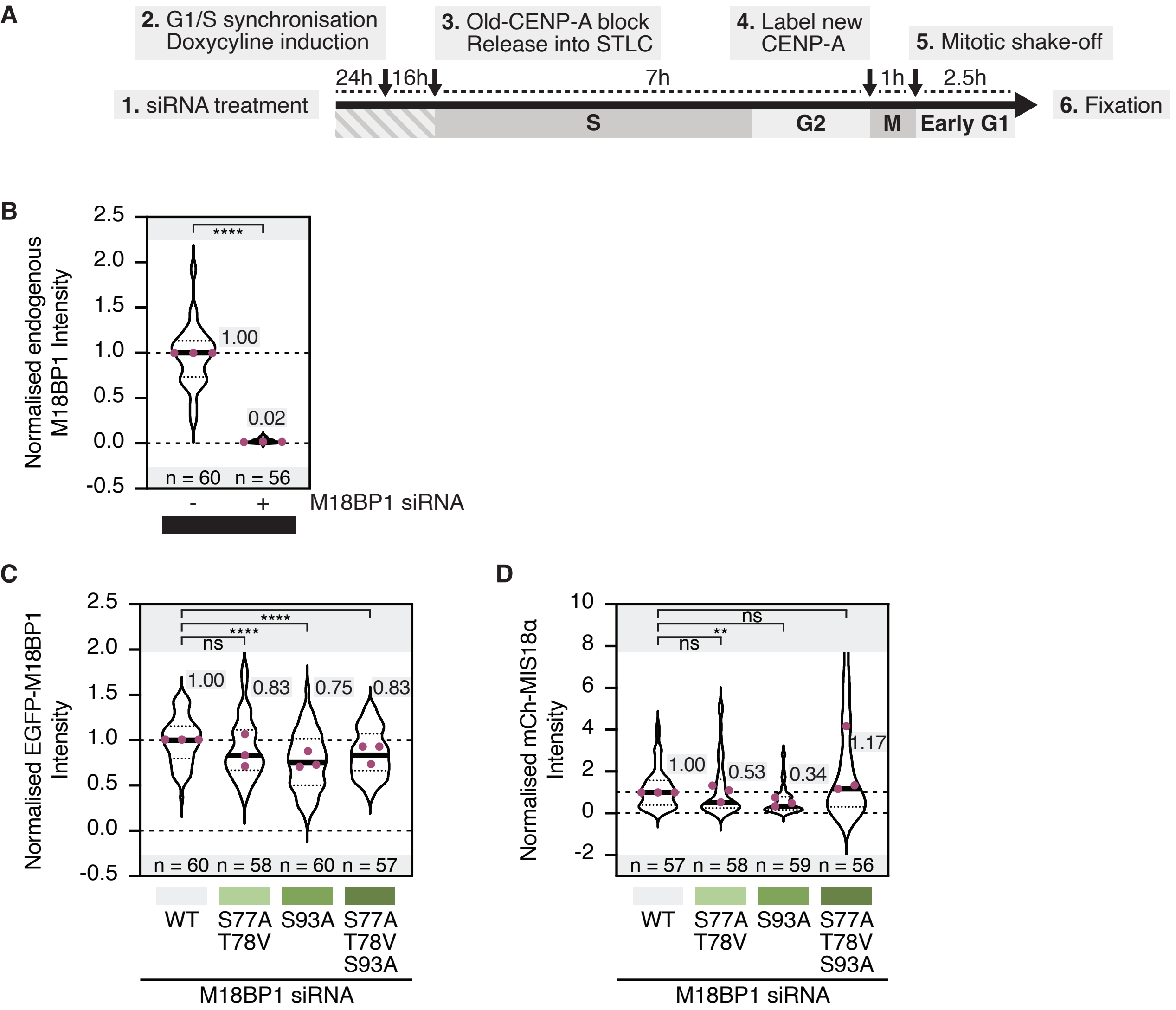

**Figure S8.** M18BP1 point mutants bind to MIS18αβ similarly to the *wild-type*.

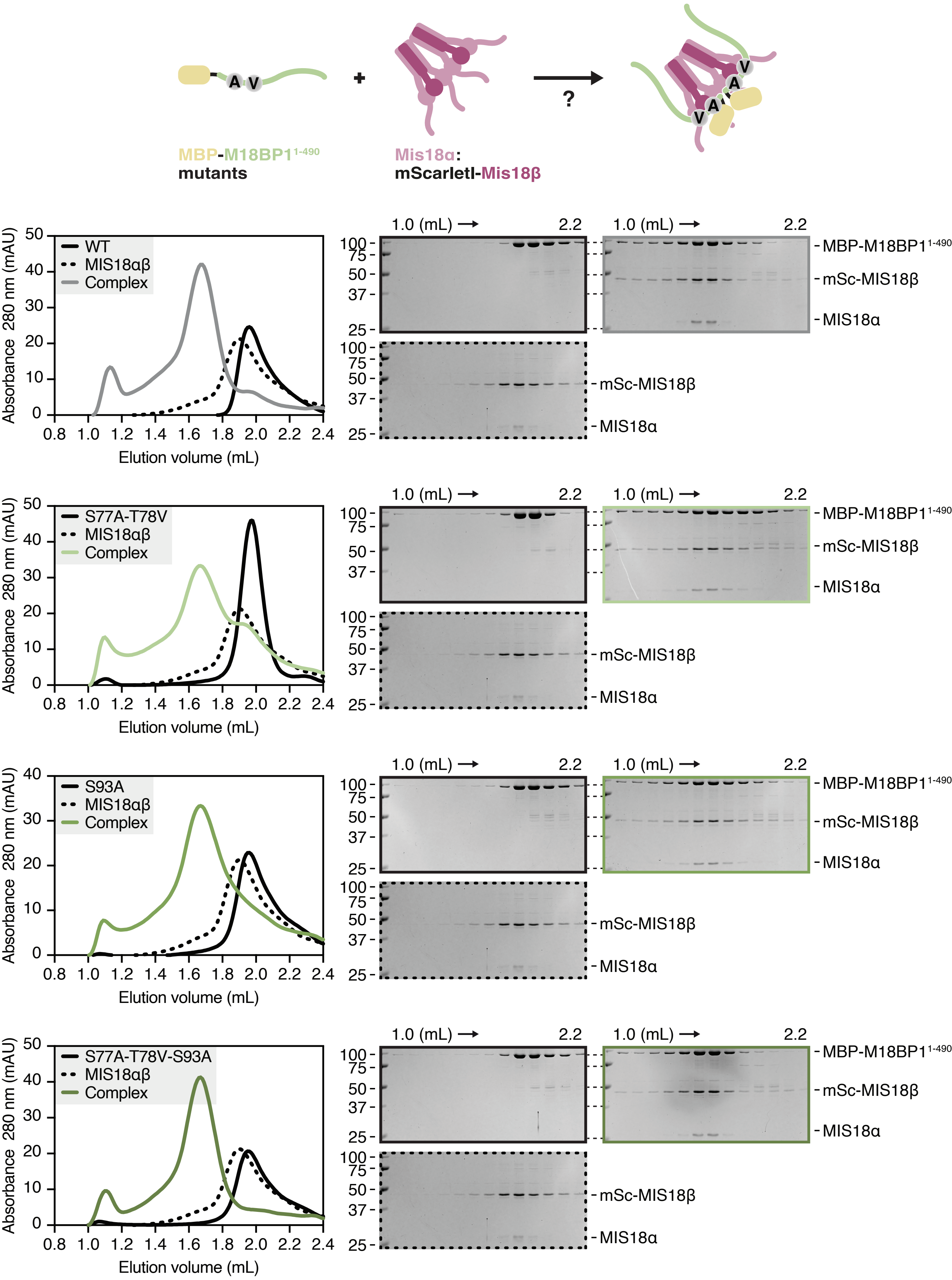

**Figure S9.** Depletion controls from Fig. 2C-D.

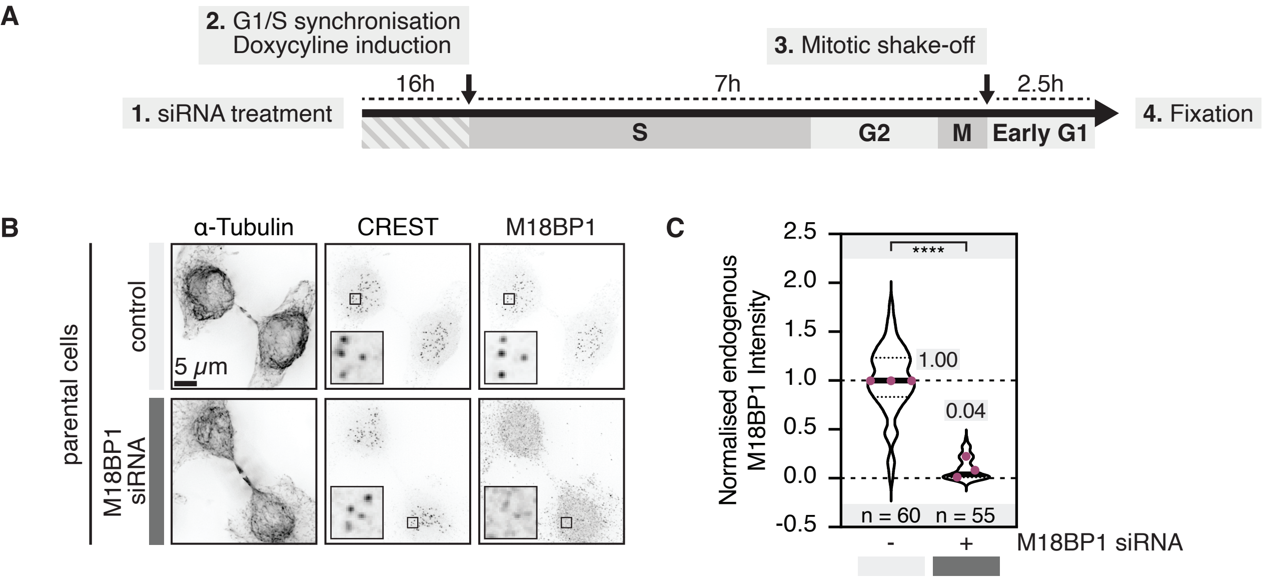

**Figure S10.** PLK1 activity and its binding to MIS18α enhances HJURP affinity for MIS18αβ.

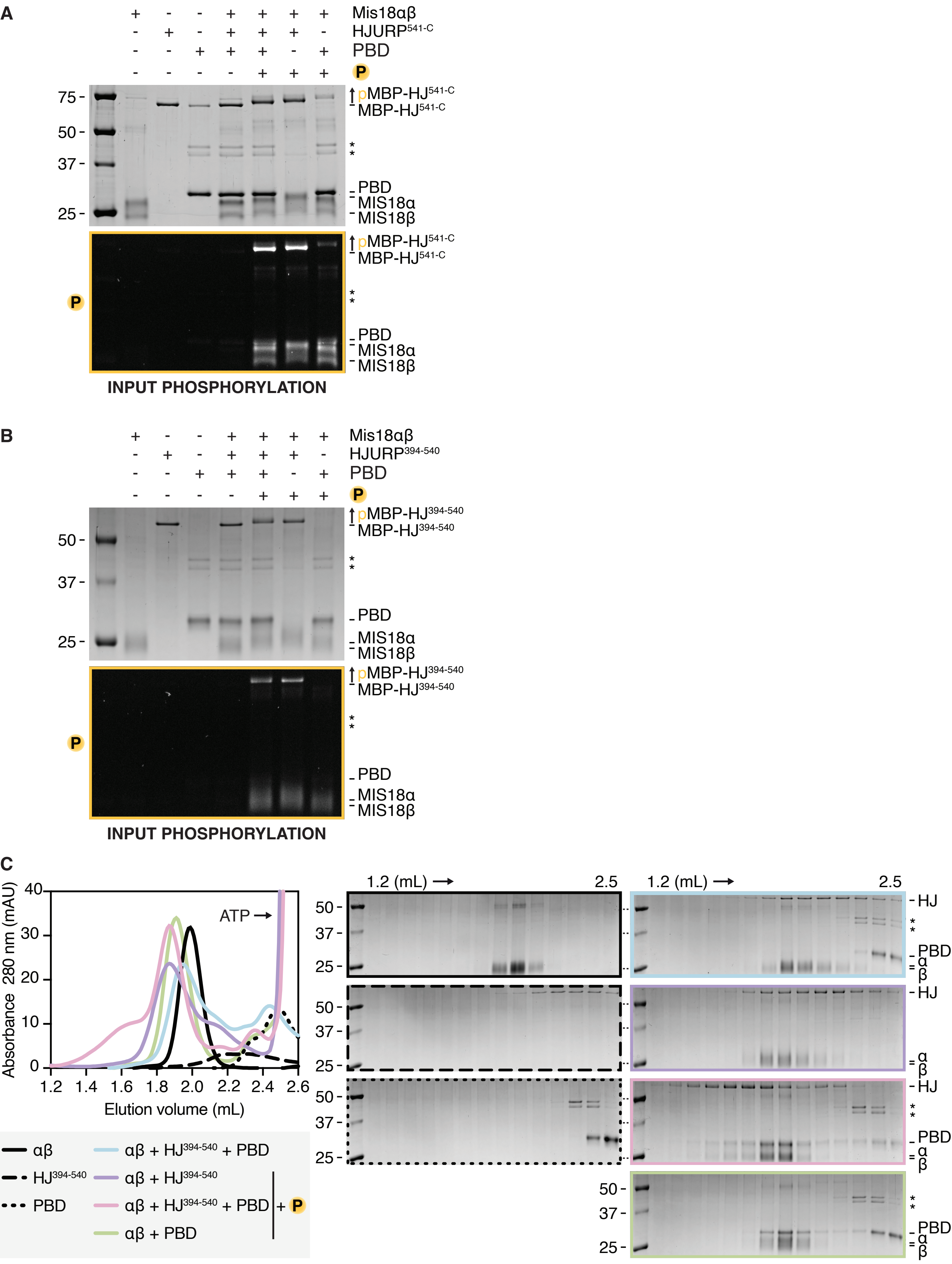

**Figure S11.** PLK1 binds to MIS18α's N-terminus in a phosphorylation-dependent manner.

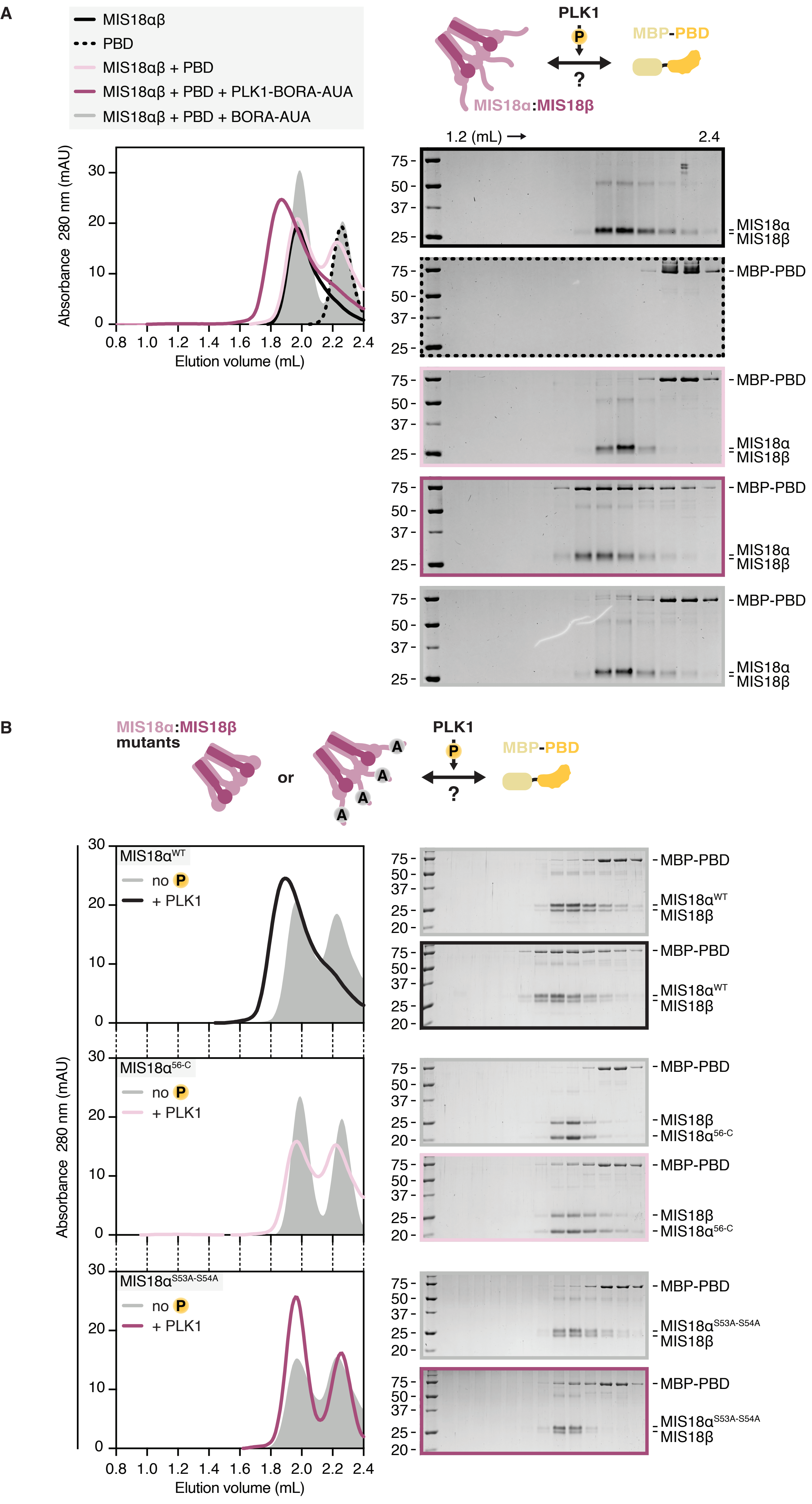

**Figure S12.** PLK1 binds to HJURP's C-terminus in a phosphorylation-dependent manner.

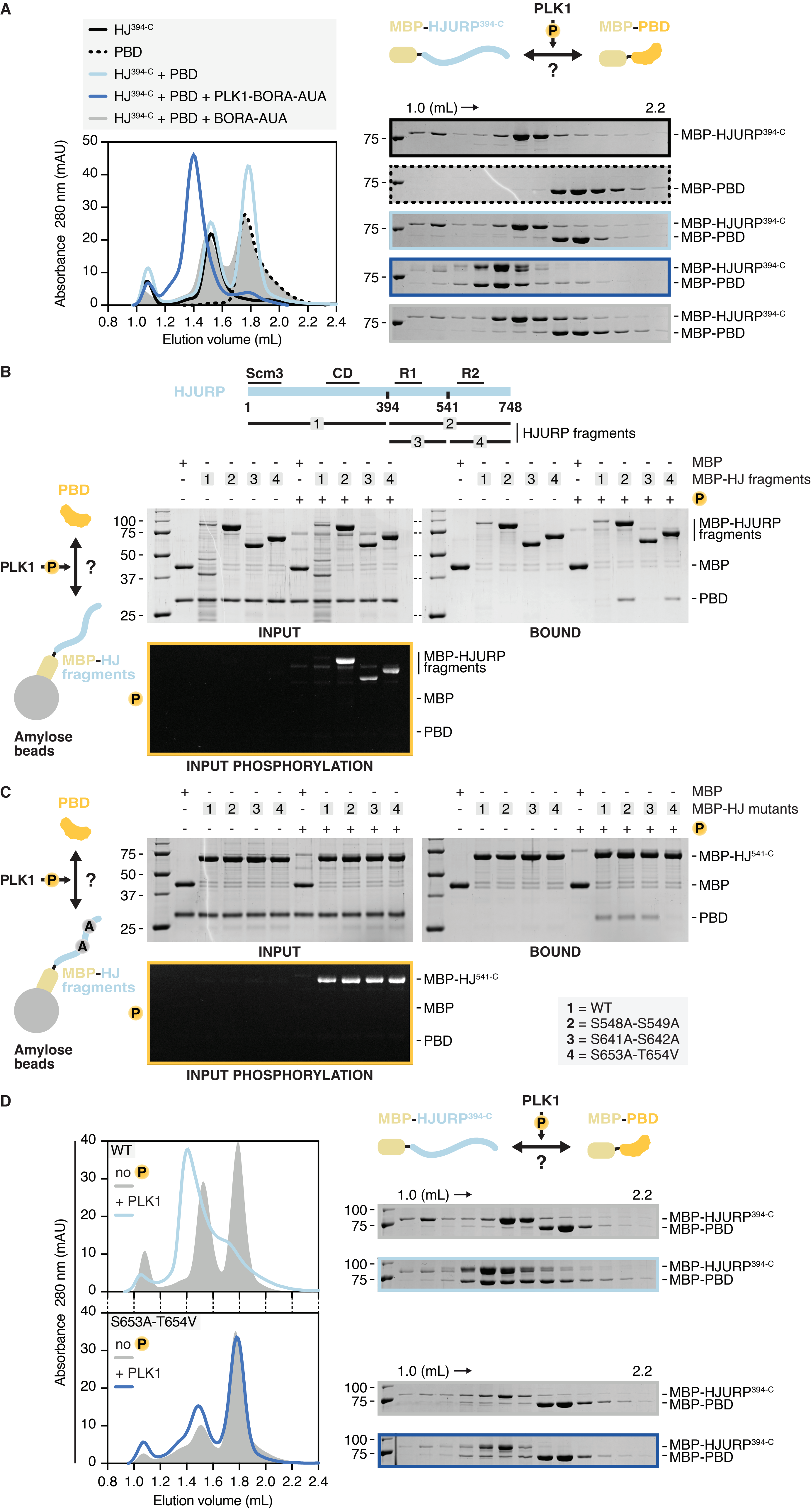

**Figure S13.** PLK1 binding to both MIS18α and HJURP is essential for the MISα18β:HJURP complex formation.

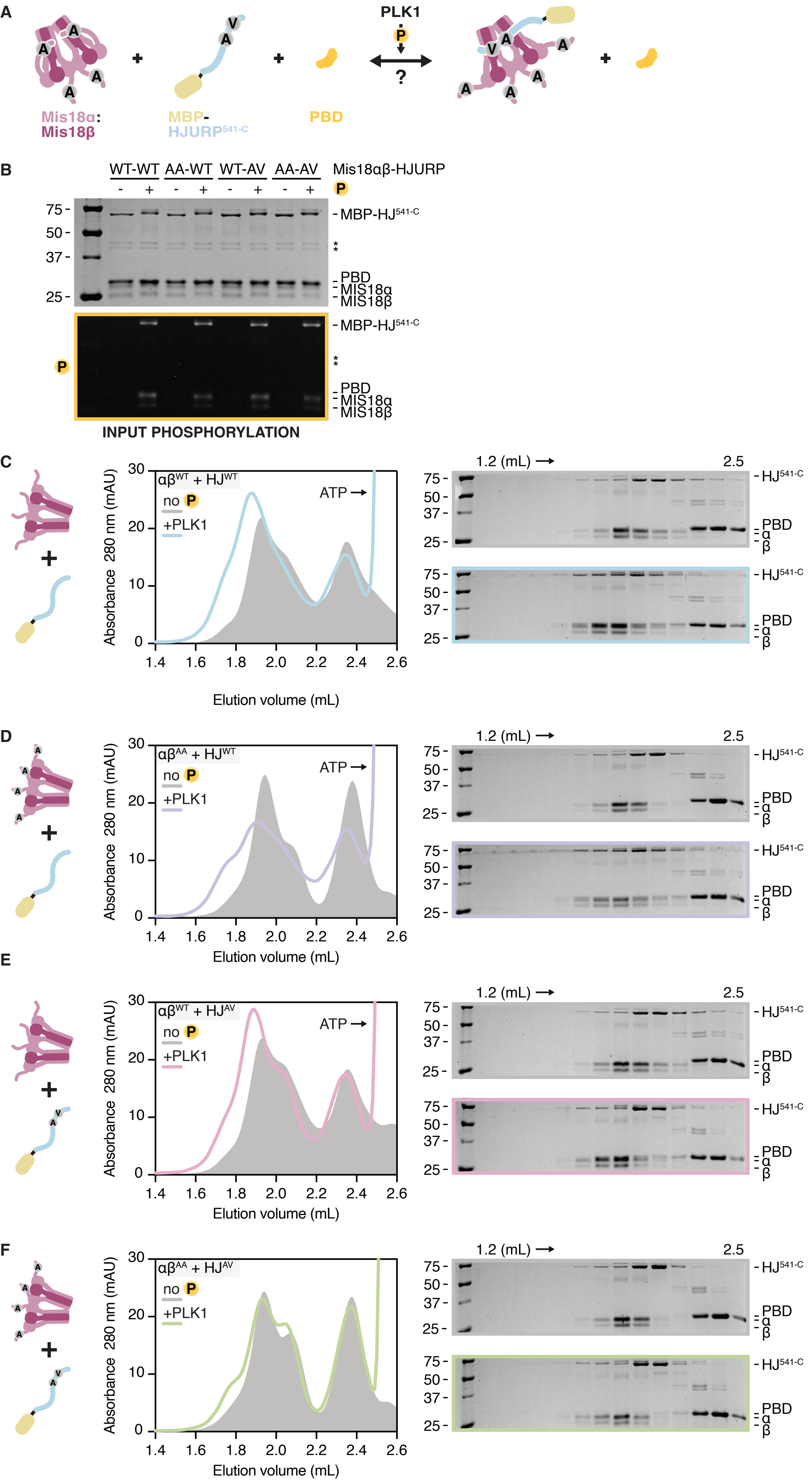

**Figure S14.** Depletion controls for Fig. 3F-H.

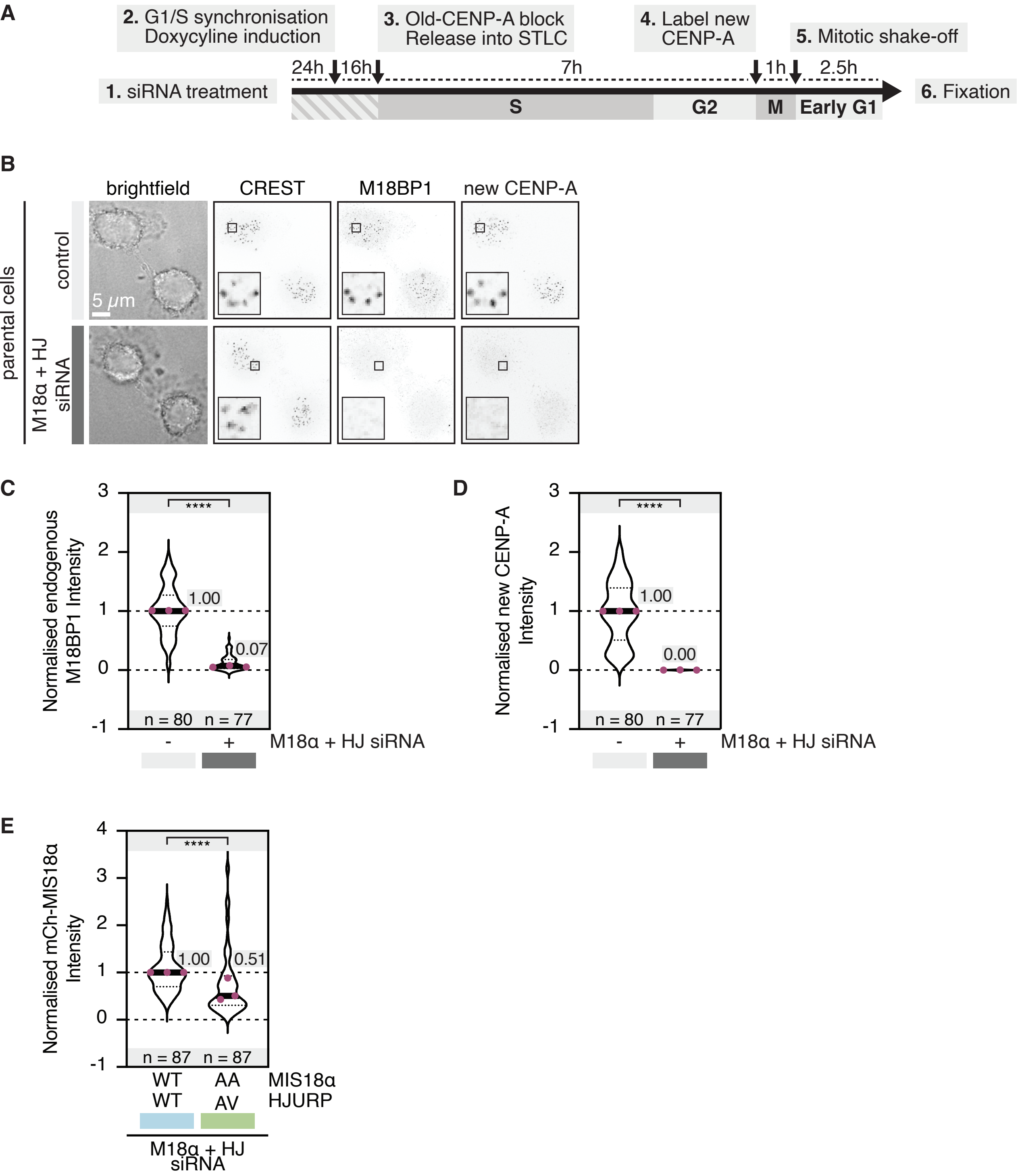

**Figure S15.** Preventing PLK1 binding to MIS18α or HJURP mildly affects new CENP-A deposition.

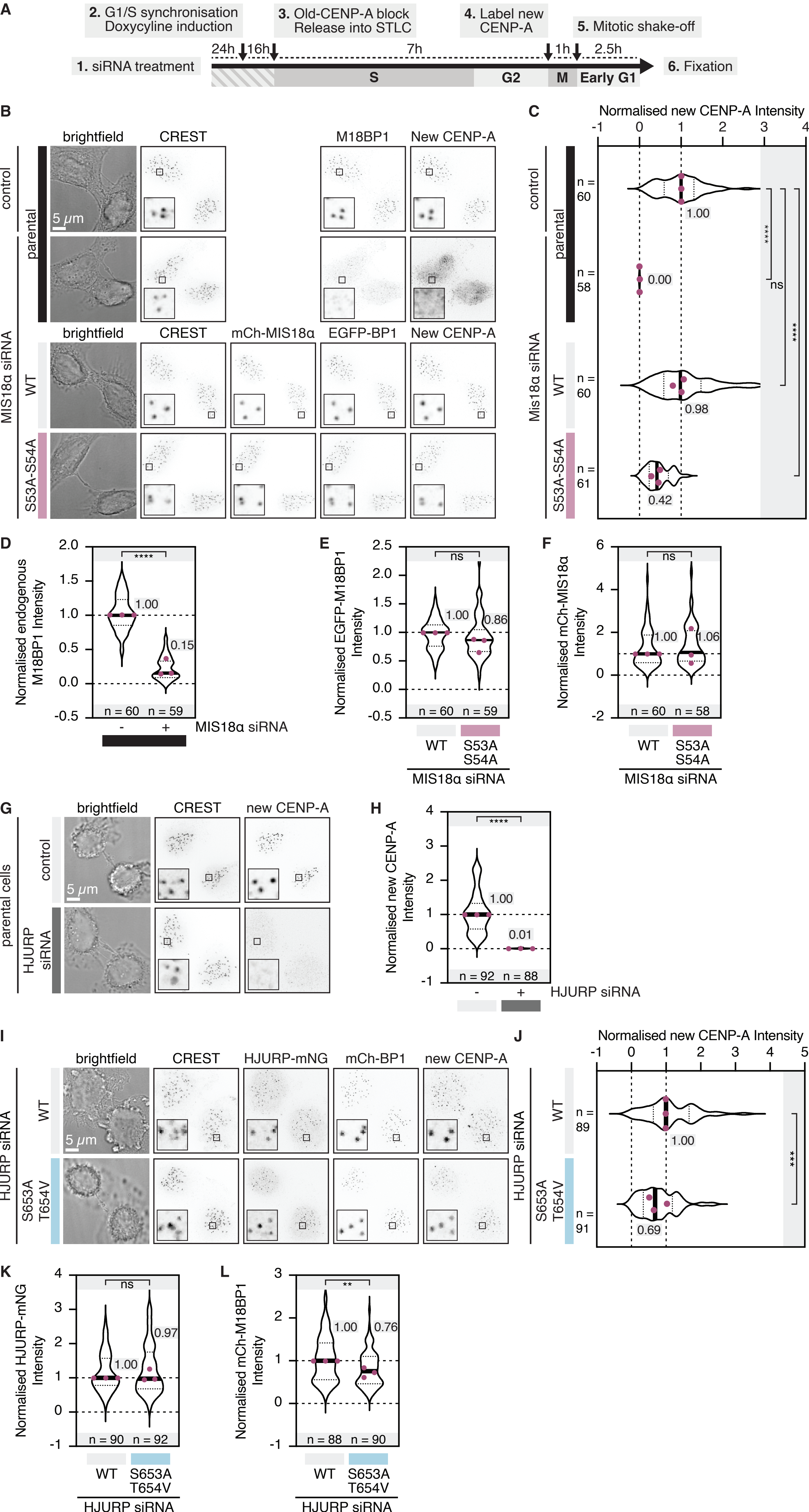

**Figure S16.** Depletion controls for Fig. 4A-B.

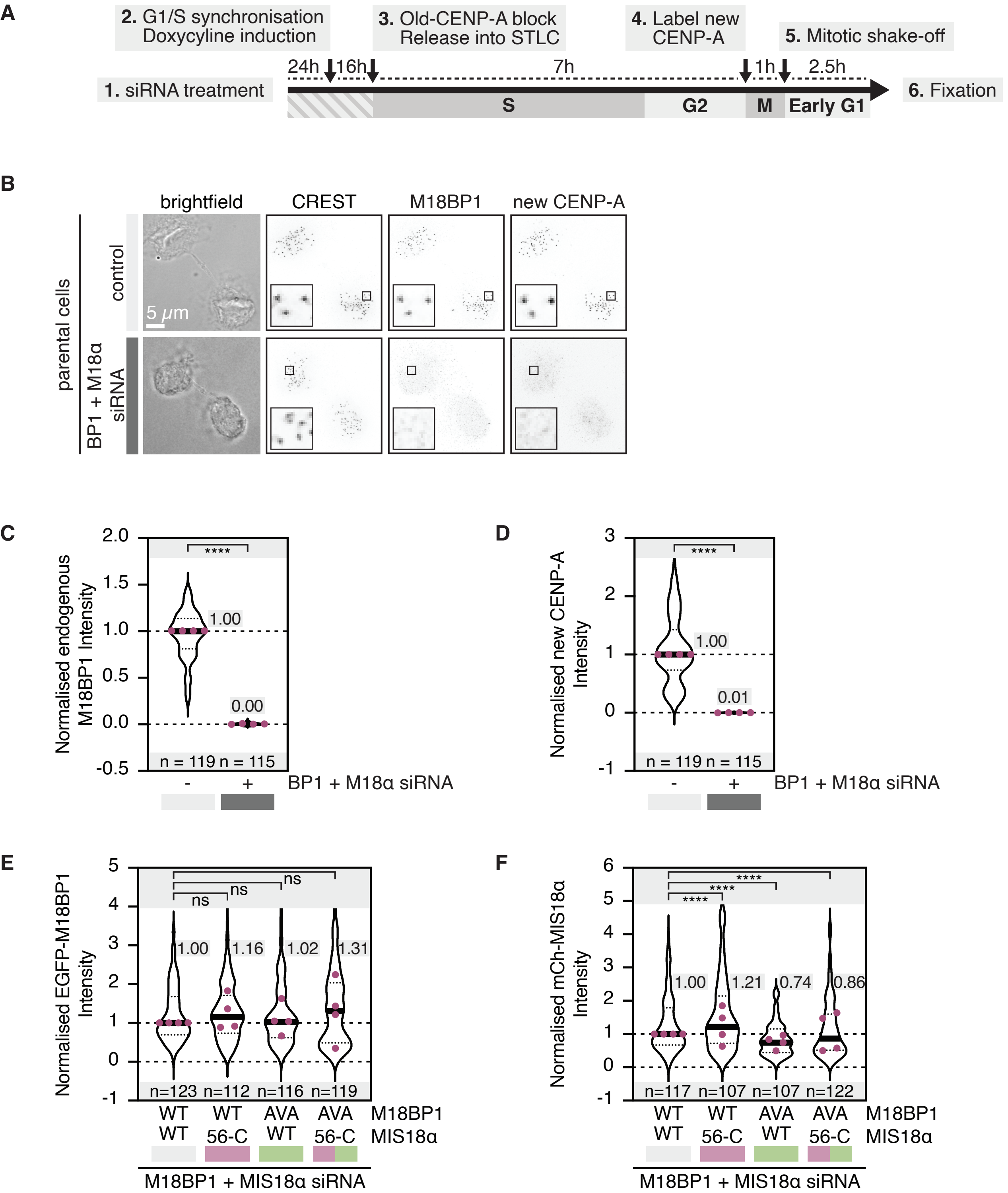

**Figure S17.** M18BP1$^{56\text{-}98}$ is a minimal PLK1-binding fragment.

A

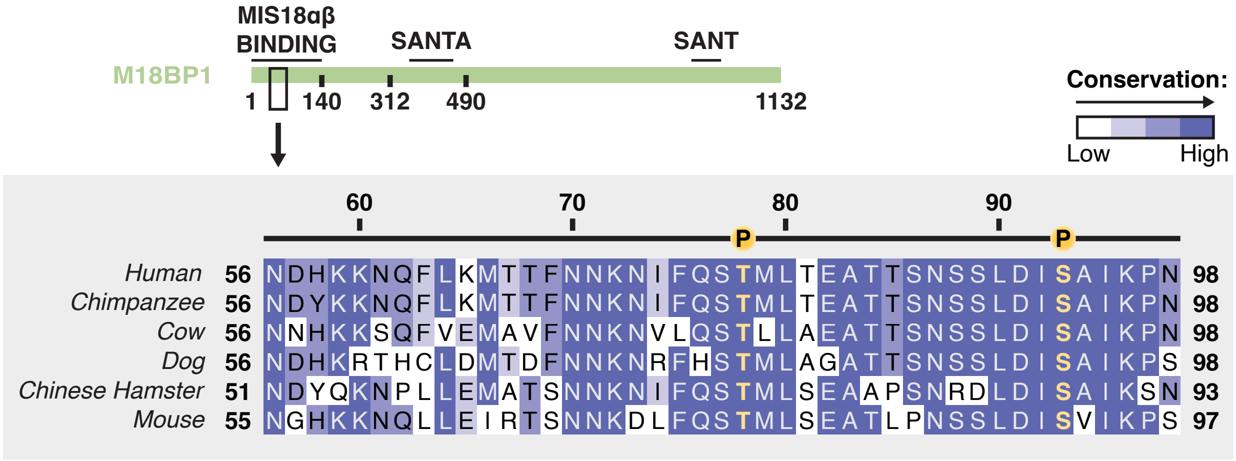


B

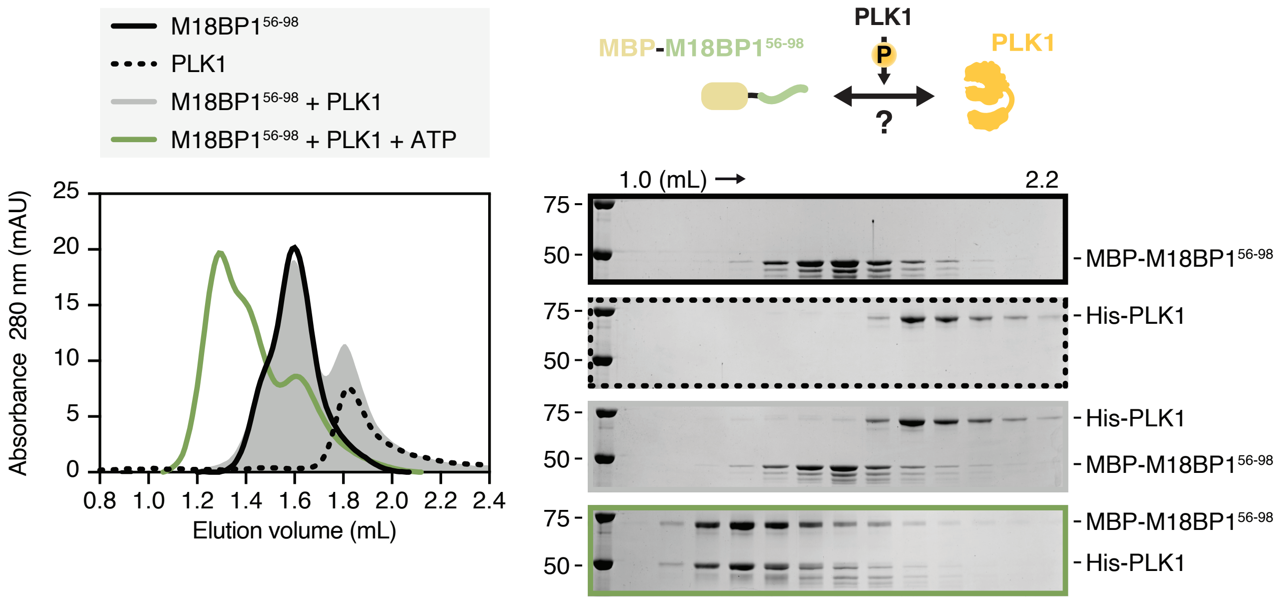

**Figure S18.** Grafting M18BP1's minimal PLK1-binding fragment onto HJURP does not sustain new CENP-A deposition in the absence of PLK1 at centromeres.

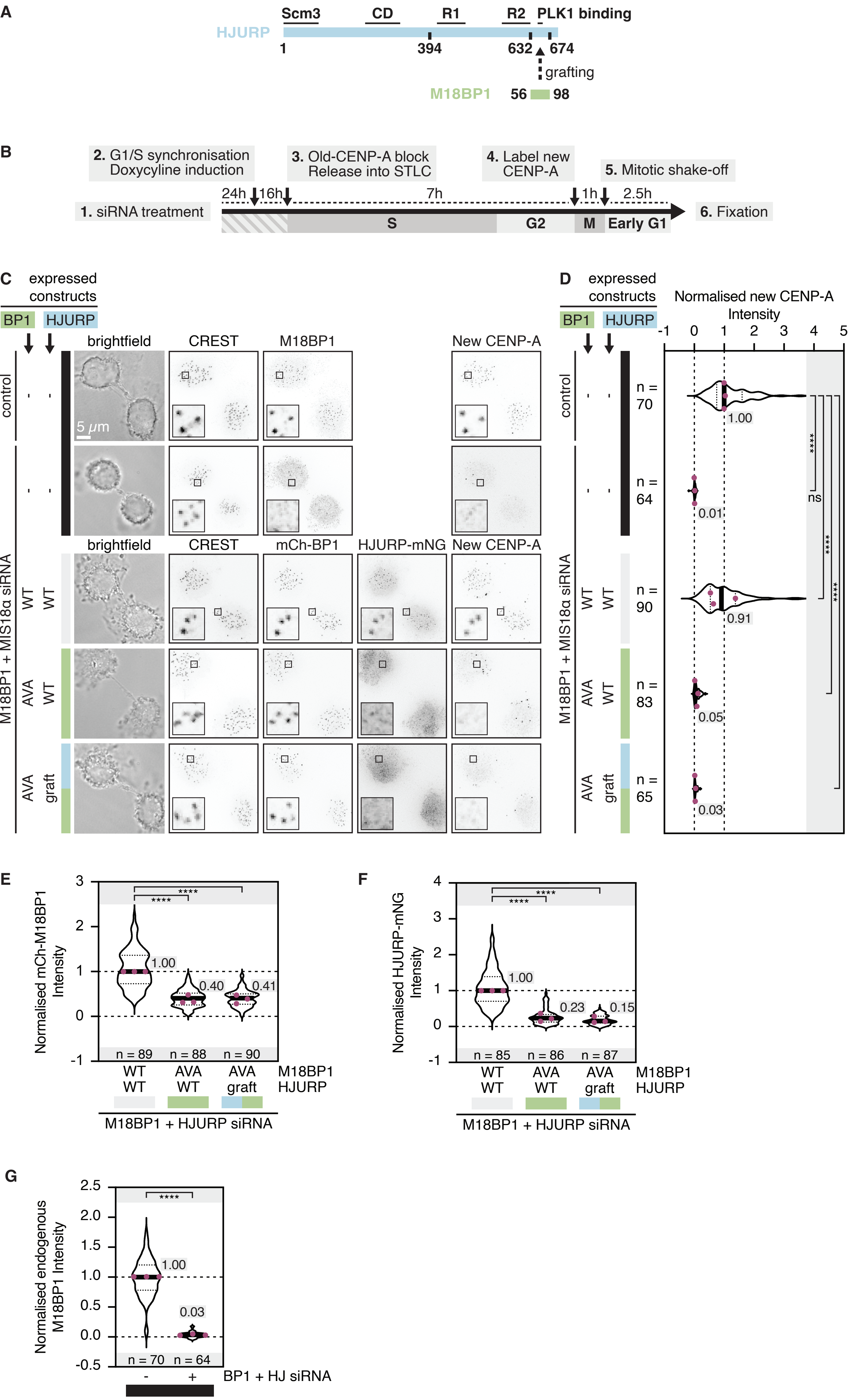

**Figure S19.** Centromeric localisation ofPLK1 and MIS18α in early G1 is not altered in cells expressing MIS18$^{S53A-S54A}$ mutants.

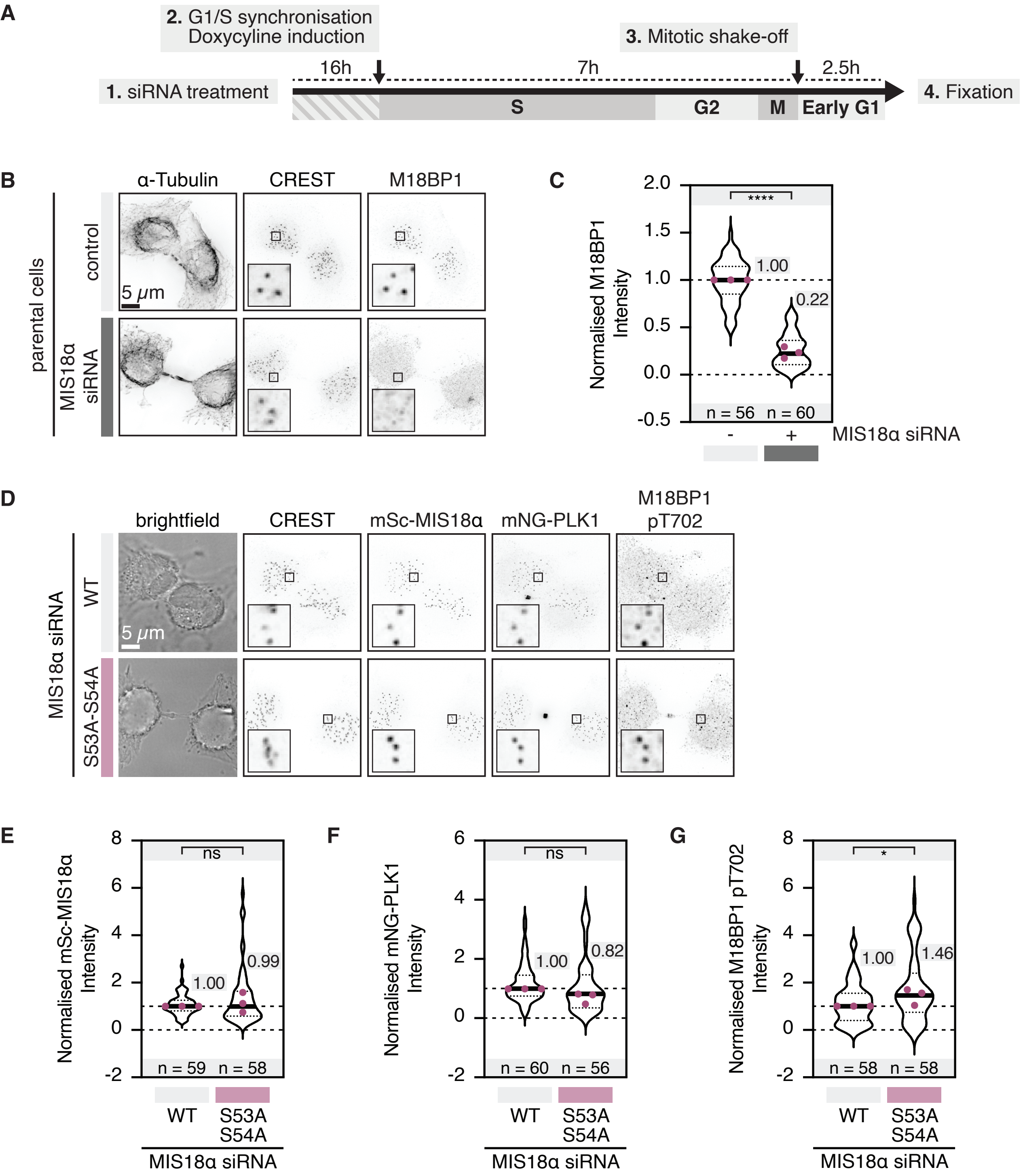

**Figure S20.** The PLK1-docking sites on M18BP1, MIS18α and HJURP are close to their respective interaction domains.

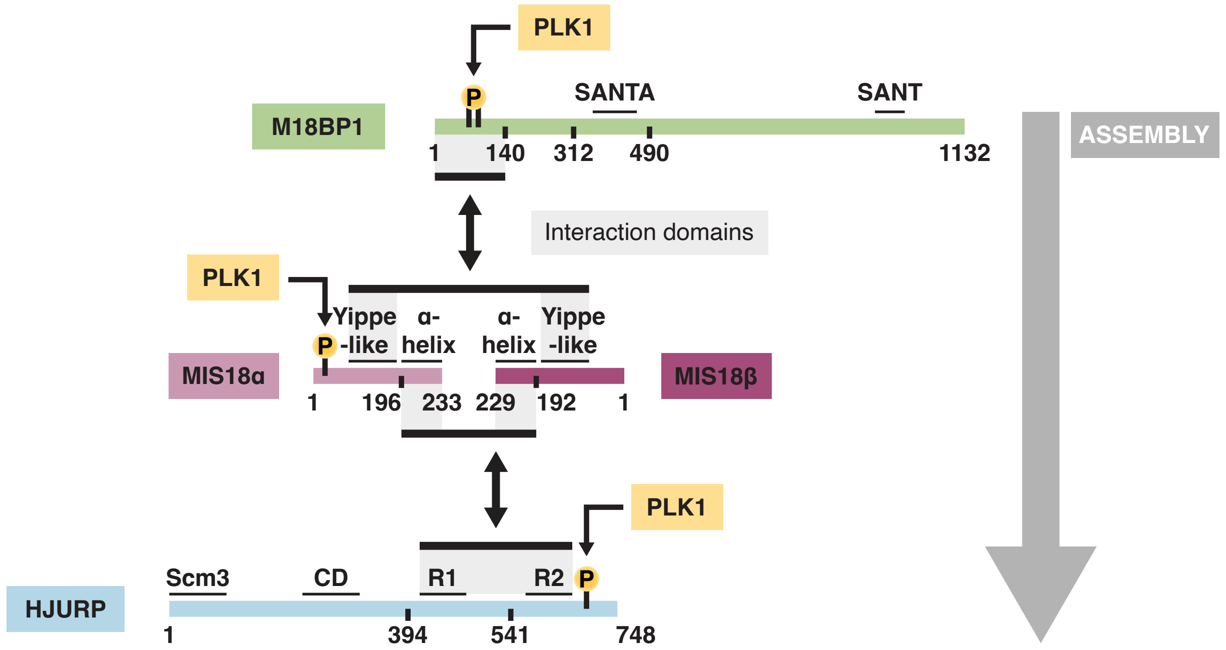
